# A high-resolution gene expression atlas of the medial and lateral domains of the gynoecium of Arabidopsis

**DOI:** 10.1101/2023.08.19.553994

**Authors:** Valentín Luna-García, Judith Jazmin Bernal Gallardo, Martin Rethoret-Pasty, Asher Pasha, Nicholas J. Provart, Stefan de Folter

## Abstract

Angiosperms are characterized by the formation of flowers, and in their inner floral whorl, one or various gynoecia are produced. These female reproductive structures are responsible for fruit and seed production, thus ensuring the reproductive competence of angiosperms. In *Arabidopsis thaliana*, the gynoecium is composed of two fused carpels with different tissues that need to develop and differentiate to consolidate a mature gynoecium and thus the reproductive competence of Arabidopsis. For these reasons, they have become the object of study for floral and fruit development. However, due to the complexity of the gynoecium, specific spatio-temporal tissues expression patterns are still scarce. In this study, we used precise laser-assisted microdissection and high-throughput RNA sequencing to describe the transcriptional profiles of the medial and lateral domain tissues of the Arabidopsis gynoecium. We provide evidence that the method used is reliable and that, in addition to corroborating gene expression patterns of previously reported regulators of these tissues, we found genes whose expression dynamics point to being involved in cytokinin and auxin homeostasis and in cell cycle progression. Furthermore, based on differential gene expression analyses, we functionally characterized several genes and found that they are involved in gynoecium development. This new resource is available via the Arabidopsis eFP browser and will serve the community in future studies on developmental and reproductive biology.

## Introduction

The gynoecium is the female reproductive structure of the flower, and it is a morphologically complex structure with a diversity of tissues and cell types. In general, upon fertilization, the gynoecium turns into a fruit, protecting the developing seeds until they are dispersed into the environment. In *Arabidopsis thaliana*, gynoecium development starts with two carpels congenitally fused to their margins, where the medial domain develops (Bowman et al., 1999; Reyes-Olalde et al., 2013; Smyth et al., 1990). At the adaxial side of the fused carpel margins, a meristematic ridge develops that is called the carpel margin meristem (CMM) (Reyes-Olalde et al., 2013; Reyes-Olalde & de Folter, 2019; Wynn et al., 2011). The lateral domain consists of the carpel wall of each carpel, which initially will grow in the shape of a hollow tube. As the gynoecium continues to develop into a mature gynoecium, it undergoes morphological changes. During the development of the medial domain, the CMM gives rise to the carpel marginal tissues, which include the placenta, ovules, septum, transmitting tract, style, and stigma, while in the lateral domains, the formation of the valve margins and valves takes place. All these tissues and structures are critical to making up a mature gynoecium ready for fertilization, where a fine action of genes and hormones is required (Balanzá et al., 2006; Bowman et al., 1999; Ferrándiz et al., 2010; Herrera-Ubaldo & de Folter, 2022; Reyes-Olalde et al., 2013; Roeder & Yanofsky, 2006; Simonini & Østergaard, 2019; Zúñiga-Mayo et al., 2019). Many years of research focused on identifying regulators involved in gynoecium development have resulted in close to a centenary of genes, many of them coding for transcription factors (Ferrándiz et al., 2010; Herrera-Ubaldo & de Folter, 2022; Reyes-Olalde et al., 2013; Roeder & Yanofsky, 2006; Simonini & Østergaard, 2019; Zúñiga-Mayo et al., 2019). Examples of medial domain expressed transcription factors are *SPATULA* (*SPT*), *HALF FILLED/CESTA* (*HAF*/*CES*), *HECATE* (*HEC1*, *HEC2,* and *HEC3*), *CUP-SHAPED COTYLEDON* (*CUC1* and *CUC2*), and *NO TRANSMITTING TRACT* (*NTT*) (Crawford et al., 2007; Crawford & Yanofsky, 2011; Gremski et al., 2007; Heisler et al., 2001; Kamiuchi et al., 2014). In the lateral domain of the gynoecium, examples of genes are: *FRUITFULL* (*FUL*), *FILAMENTOUS FLOWER* (*FIL*), *JAGGED* (*JAG*), and *CRABS CLAW* (*CRC*) (Bowman & Smyth, 1999; Dinneny et al., 2005; Gu et al., 1998). In addition, experimental evidence of expression patterns of reporters and genes (such as those involved in biosynthesis, transport, and transcriptional response) related to cytokinin and auxin pathways (e.g., Marsch-Martínez et al., 2012b; Moubayidin & Østergaard, 2014; Müller et al., 2017; Reyes-Olalde et al., 2017) indicates that hormones are playing a key role during carpel marginal tissue development. Knowledge of all these regulators represents a major advance in the study of the molecular control of gynoecium development. However, developing tissue-specific expression profiles could help elucidate all spatial regulatory events, ultimately uncovering biological pathways and regulatory networks that have been undetectable during development (Belmonte et al., 2013; Taylor-Teeples et al., 2011).

In the last few years, techniques such as translating ribosome affinity purification (TRAP), isolation of nuclei tagged in specific cell types (INTACT) and Fluorescent Activated Cell Sorting (FACS) have been used to isolate and characterize transcriptomic profiles of specific cell populations (Carter et al., 2013; Deal & Henikoff, 2011; Zanetti et al., 2005). In addition to the need to create transgenic or specific marker lines to use these techniques, not all tissues are suitable starting materials for the production of protoplasts. More recent techniques are single cell and single nuclei sequencing (Cervantes-Pérez et al., 2022; Denyer & Timmermans, 2022; Rich-Griffin et al., 2020), though these techniques are still challenging, and the physical location of the cells is lost. Gene expression patterns are predicted based on clustering strategies. When no marker genes are present in a cluster, it is hard or impossible to predict the identity of the original cell population. Therefore, creating transcriptome atlases of specific tissues and cell types during plant development is very useful for future studies with the mentioned techniques. One technique to isolate specific cell types from plant tissues and organs, is Laser-Assisted Microdissection (LAM) microscopy. This technique has been consolidated for collecting cell populations from their tissue context, regardless of their complexity (Chávez Montes et al., 2016; Day et al., 2005; Florez Rueda et al., 2016; Kerk et al., 2003; Wuest & Grossniklaus, 2014). Despite the fact that the LAM technique is also labor intensive, the advantage is, unlike other techniques, that it does not require the generation of marker lines, protoplasts, or the need for previous information on a large number of marker genes, except for knowing the morphology of the tissues to be collected, resulting in a large number of marker genes (Chávez Montes et al., 2016; Day et al., 2005; Florez Rueda et al., 2016; Wuest & Grossniklaus, 2014).

A technique known as LAM coupled with RNA-Seq was used here to provide high spatial and temporal resolution for the expression of specific transcriptional programs during the development of the carpel margin meristem (CMM), carpel walls (PC), septum (SEP), and valves (VV) in Arabidopsis. Our analyses classified around 14,000 to 15,000 expressed genes in the specific cell types. Our data is also available in the Arabidopsis eFP Browser for easy data exploration and hypothesis-making. We describe the expression dynamics of transcription factors, the known gynoecium regulators, auxin and cytokinin pathway genes, and the cell cycle genes. Furthermore, around 1500 to 4000 differentially expressed genes (DEGs) for each tissue were detected. We selected 7 DEGs for the medial tissues (CMM and SEP) and using T-DNA insertion lines for functional analyses, found their involvement in reproductive development in Arabidopsis.

## Results

### Laser-assisted microdissection and RNA-seq

With the goal of elucidating which genes are tissue-specifically expressed during the development of the gynoecium, we used the laser-assisted microdissection (LAM) method (Chávez Montes et al., 2016) combined with RNA-Seq. We generated a high-resolution expression map by collecting different tissues at two time points. Stage 7 gynoecia tissues included the carpel margin meristem (CMM) and the young carpel wall or presumptive carpel wall (PC). Later in development, at stage 10 gynoecia, the septum (SEP) and valve (VV) tissues were collected (Figure 1). The selection of these tissues allows us to discover gene expression in the medial and lateral tissues during gynoecium development and observe how gene expression patterns change from tissues with meristematic activity (CMM) or not much-differentiated tissue yet (PC) to tissues that likely lost meristematic activity (SEP) and are differentiated (SEP and VV), respectively.

**Figure 1.**
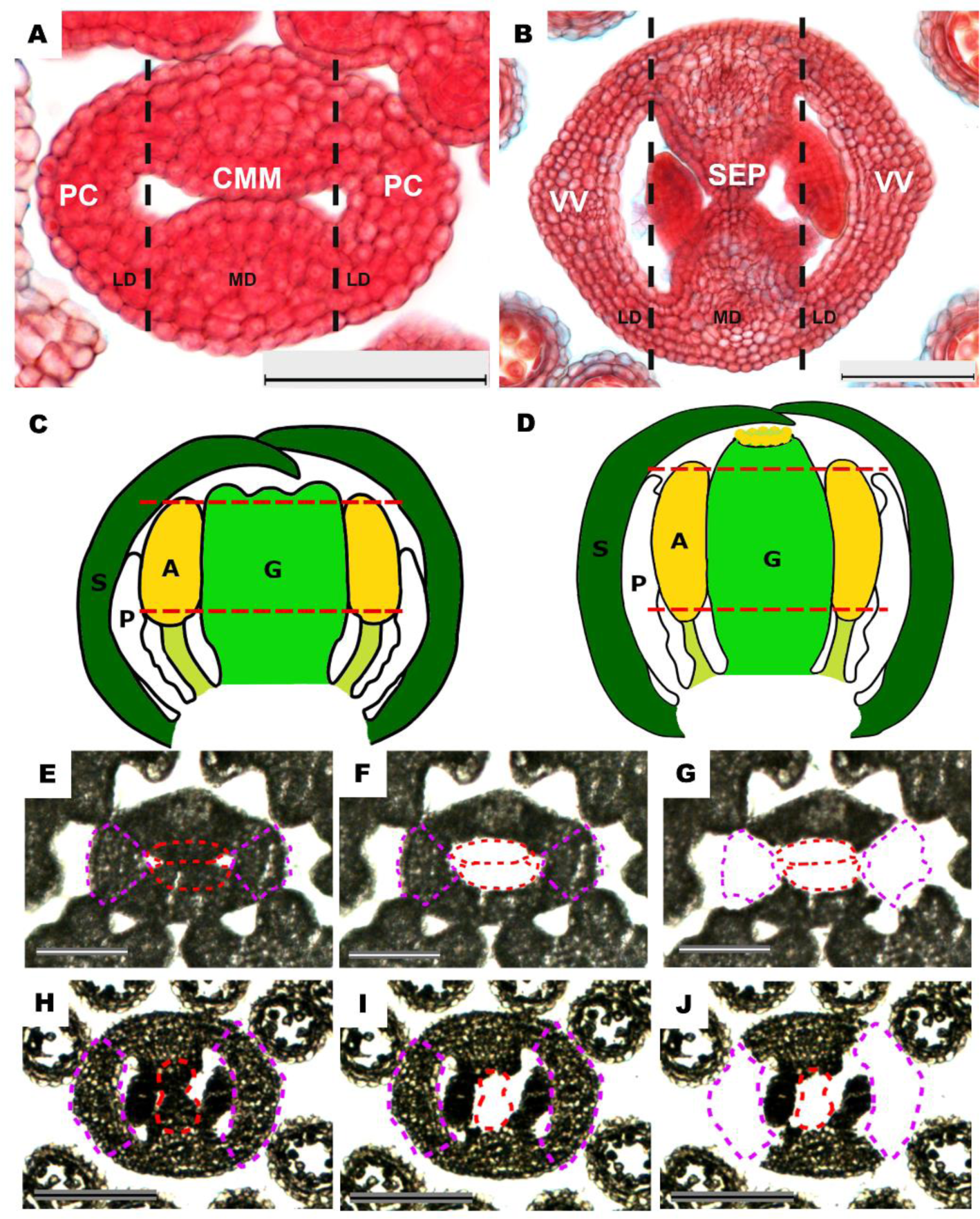
Laser-assisted microdissection. **A and B)** Cross-sections of *Arabidopsis thaliana* gynoecia stained with neutral red and alcian blue at stage 7 and 10, respectively. Dashed black lines separate the medial (MD) and lateral (LD) domains. CMM – carpel margin meristem, PC – presumptive carpel wall, SEP – septum and VV – valves. Scale bars are 20 µm. **C and D)** Drawings of longitudinal sections of a flower in stage 7 and 10, respectively. Dashed red lines show the part of the gynoecium used to collect the CMM and PC tissues (C), and SEP and VV tissues (D). S – sepal, P – petal, A – stamen, G – gynoecium. **E to J)** Cross-sections of gynoecia at stage 7 (E-G) and at stage 10 (H-J). **F, G, I, J)** Cross-sections after the collection of the tissue of interest: CMM (F), PC (G), SEP (I), and VV (J) tissues. E-J: Scale bars are 20 µm.

First, transverse sections were made of around 240 gynoecia on a microtome and mounted on slides, followed by laser microdissection to collect each specific tissue, and extract RNA from each tissue in triplicate (see M&M for details and Supplemental Figure 1). A total RNA yield of 3.3 to 12.3 ng/µL with an OD260/280 of 1.7 ± 0.2 and an OD260/230 of 1.0 ± 0.2 was obtained (Supplemental Table S1). We used 10 ng of RNA input per sample for cDNA synthesis using the SMART-Seq v4 Ultra Low Input RNA Kit, followed by library preparations (see M&M for details).

The sequencing was performed in one lane of a HiSeq2000 Illumina to generate paired-end libraries with an average of 43 million reads (150 pb) after filtering adapters and low-quality reads. On average, 81% of the reads aligned to the *Arabidopsis thaliana* Araport11 reference transcriptome, while the remaining reads aligned to the rRNA gene (Supplemental Table S1).

For exploratory analysis of the results of the 12 libraries, normalized reads of all samples were used to perform a principal component analysis (PCA). The PCA illustrated that the two primary PC axes explain 75% of the total variance among all samples, as well as four clusters of the four tissue types with their respective replicates (Supplemental Figure 2), indicating that samples of the four tissue types are different and that the replicates are uniform.

### Gene Expression Atlas Analysis

To generate a gene expression atlas, we first generated a list of all Arabidopsis genes and their detected expression in each tissue in TPM (Transcripts Per Kilobase Million) (Supplemental Table S2). Secondly, lowly expressed genes were filtered using a cut-off of ≥4 TPM, which resulted in 15059, 14251, 14318, and 13864 genes expressed in the CMM, PC, SEP, and VV tissues, respectively (Figure 2A; Supplemental Tables S3-S6). To make this data easily available as a resource for the community, all expression data is searchable in the Arabidopsis eFP browser (Winter et al., 2007); examples of the eFP browser results are given below (Figures 4D, 6A, 7A, and G).

**Figure 2.**
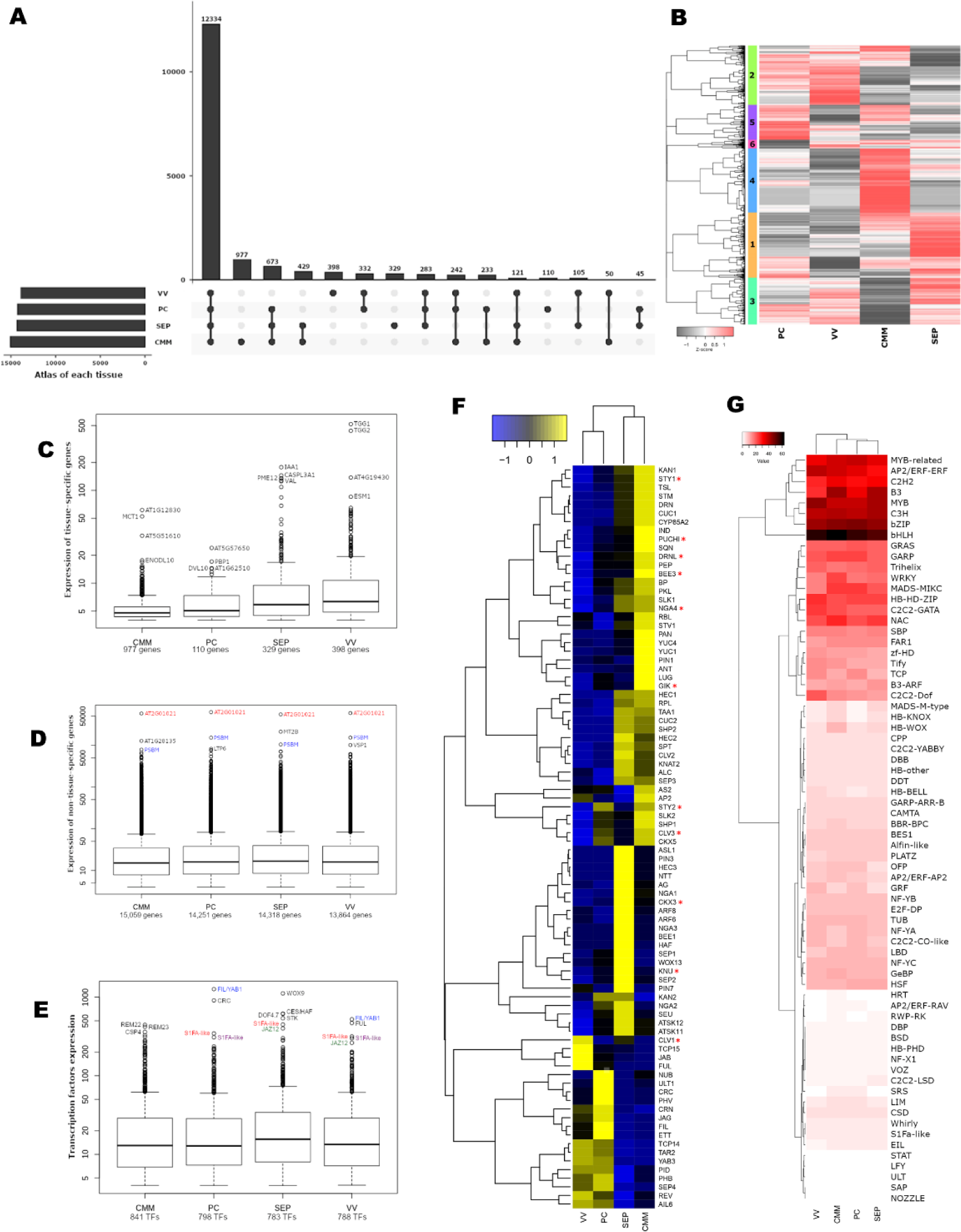
Gene expression atlas of medial and lateral domain tissue analyses. **A)** Upset plot of the number of expressed genes in each tissue, showing the intersection of shared genes. **B)** k-means clustering of the gene expression dynamics of each tissue (CMM, PC, SEP, and VV). **C)** Expression level of tissue-specific genes in each tissue. **D)** Expression level of non-tissue-specific genes (≥ 4 TPM) in each tissue. **E)** Expression level of transcription factors in each tissue. **F)** Heatmap of the expression profile of the 86 genes involved in medial and lateral domain tissue development (Reyes-Olalde et al. 2013). * Indicates genes with very low expression (< 4 TPM). **G)** Representation of the number of transcription factors expressed in each tissue, grouped by transcription factor family.

Next, we explored gene expression profiles by performing a K-means clustering approach, which resulted in six clusters presented as a heatmap in Figure 2B. The number of genes present in each cluster (Supplemental Table S7), is as follows: Cluster 1 contains 3895 genes, which are highly expressed in SEP tissue; Cluster 2 contains 3540 genes, of which the majority have a higher expression in VV tissue; Cluster 4 contains 3819 genes that are highly expressed in CMM tissue; Cluster 5 contains 2089 genes of which the majority are highly expressed in PC tissue; and the remaining two clusters, Clusters 3 and 6, are characterized by genes whose expression is shared in two or three tissues.

Clusters with genes highly expressed in one of the four tissues, are not automatically related to genes uniquely expressed in that tissue (Figure 2B). The number of tissue-specific expressed genes (≥4 TPM) for CMM tissue is 977, for PC tissue it is 110, for SEP tissue it is 329, and for VV tissue it is 398 (Figure 2A, Supplemental Table S8). Of these tissue-specific genes, the number of tissue-specific transcription factors for each tissue is the following: 67 for CMM, 17 for PC, 30 for SEP, and 44 for VV (Supplemental Table S9). Related to gene expression levels, most of the tissue-specific genes have an expression level of around 5 to 6 TPM (Figure 2C, Supplemental Table S10), while most non-tissue-specific genes are expressed around 15 to 17 TPM, in addition to a group of genes with much higher expression (Figure 2D, Supplemental Tables S3-S6). Furthermore, when taking into account only the transcription factors of the non-tissue-specific genes, expression levels are also around 15 to 17 TPM, in addition to a group of genes with higher levels (Figure 2E, Supplemental Table S11). In each boxplot, the highest-expressed genes are indicated.

### Expression analysis of known gynoecium regulators

Based on a review, at least 86 genes are involved in gynoecium development in Arabidopsis. Many genes encode transcription factors, but also transcriptional co-regulators, hormonal pathway components, and other genes with important functions (Reyes-Olalde et al., 2013). All genes were detected, though some had very low expression (indicated with an asterisk in Figure 2F; Supplemental Table S12). After applying a cut-off value of ≥4 TPM, 75 out of 86 genes were considered to be expressed in one or more of the four tissues (Figure 2F).

We observed that most of the genes (66 of 86 genes; 77%) are expressed in the medial domain (CMM and PC). Furthermore, many genes are preferentially expressed in only one tissue, and less in other tissues. Interestingly, only three real tissue-specific genes were detected, which all have a link to the hormone auxin. Two genes are specific for CMM tissue: the auxin biosynthesis genes *YUCCA 1* (*YUC1*) and *DORNRÖSCHEN* (*DRN*), related to floral meristem identity and organ initiation, which is a direct target of MONOPTEROS (MP/ARF5). The third specific gene is the auxin efflux transporter *PIN-FORMED 3* (*PIN3*) in the septum.

In general, the observed gene expression patterns are related to the functions of the genes (Figure 2F; Reyes-Olalde et al., 2013). In the CMM tissue, transcription factors such as *SHOOT MERISTEMLESS* (*STM*), *BREVIPEDICELLUS* (*BP*), *PERIANTHIA* (*PAN*), *CUCs*, *AINTEGUMENTA* (*ANT*), and *SPT* are related to meristem activity. Transcription factors important for septum development present in SEP tissue are *NTT*, *HECs*, *BR-ENHANCED EXPRESSION 1* (*BEE1*), *HAF*, *SPT*, and *AUXIN RESPONSE FACTOR 6* and *8* (*ARF6* and *ARF8*). Among the transcription factors related to carpel wall and valve development in the lateral domain (PC and VV), we find *FUL*, *CRC*, *JAG*, *FIL*, and *ETTIN* (*ETT*/*ARF3*).

Lastly, as mentioned above, 11 out of 86 genes have very low expression in the four tissues (CMM, PC, SEP, and VV) analyzed, lower than 4 TPM (indicated with a gray color in Supplemental Table S12), though they are important for gynoecium development. These genes are the following: *BEE3*, *DORNRÖSCHEN-LIKE* (*DRNL*), *CLAVATA 1* (*CLV1*), *CLAVATA 3* (*CLV3*), *GIANT KILLER* (*GIK*), *STYLISH 1* (*STY1*), *NGATHA 4* (*NGA4*), *STYLISH 2* (*STY2*), *KNUCKLE* (*KNU)*, *PUCHI*, and *CYTOKININ OXIDASE 3* (*CKX3*). These 11 genes are not considered in our analyses.

### The involvement of transcription factor family members in gynoecium development

Many of the known regulators of gynoecium development are transcription factors (Figure 2F). Based on the iTak database (Zheng et al., 2016), Arabidopsis has around 1700 transcription factors. Around 45% of these 1700 transcription factors are expressed in each of the tissues, belonging to ∼68 of the total 70 classified transcription factor families (Supplemental Tables S11 and S13).

As illustrated in Figure 2G, a large number of expressed transcription factors belong to the basic/helix-loop-helix (bHLH) and basic region/leucine zipper (bZIP) families, two families that have been described as controlling a diversity of plant and floral developmental processes. The results suggest that these members are the main regulators of tissue development in both the medial and lateral domains. On the other hand, the results suggest that many members of the B3 transcription factor family are regulating the development of the medial domain tissues (CMM and SEP), a superfamily that encompasses well-characterized families such as the Auxin Response Factor (ARF) and Reproductive Meristem (REM) families, which are participating in the establishment of the cell lineage of this domain. Meanwhile, the lateral domain is characterized by the presence of many members of the MYB and MYB-related transcription factor families, which have been described as crucial to controlling proliferation and differentiation in several cell types. Furthermore, many members of the AP2 and C2H2 type Zinc finger transcription factor families are seen in the different tissues, which are mainly involved in plant growth and development. Lastly to mention, quite a large number of members belonging to the MADS domain family are observed, well known for their involvement in floral organ development, including the gynoecium.

### Auxin and cytokinin homeostasis in the medial and lateral domains of the gynoecium

Plant hormones, especially auxin and cytokinin, are important for gynoecium development (Deb et al., 2018; Herrera-Ubaldo & de Folter, 2022; Larsson et al., 2013; Marsch-Martínez et al., 2012a; Marsch-Martínez & de Folter, 2016; Robert et al., 2015; Sehra & Franks, 2015). It is known that hormone homeostasis is a complex process that involves hormone biosynthesis, transport, response, and degradation pathways. Focusing on auxin and cytokinin, these two plant hormones have been referred to as the “Yin-Yang” of plant development (Schaller et al., 2015), because although these two plant hormones have opposite functions, they act together for proper plant and flower development.

To understand the correlation between auxin and cytokinin pathway genes during medial and lateral domain development, we analyzed the expression of 108 and 78 genes with annotated functions during auxin and cytokinin homeostasis mechanisms, respectively (Figures 3A, B, and Supplemental Tables S14-S15). Of the 108 auxin pathway-related genes, 59 were expressed (≥ 4 TPM) and between 25 and 31 were deferentially expressed in one of the tissues (considering an FDR< 0.05), while for the 78 cytokinin pathway-related genes, 44 were expressed (≥ 4 TPM) and between 11 and 32 were deferentially expressed in one of the tissues (considering an FDR< 0.05) (Figures 3A and B).

**Figure 3.**
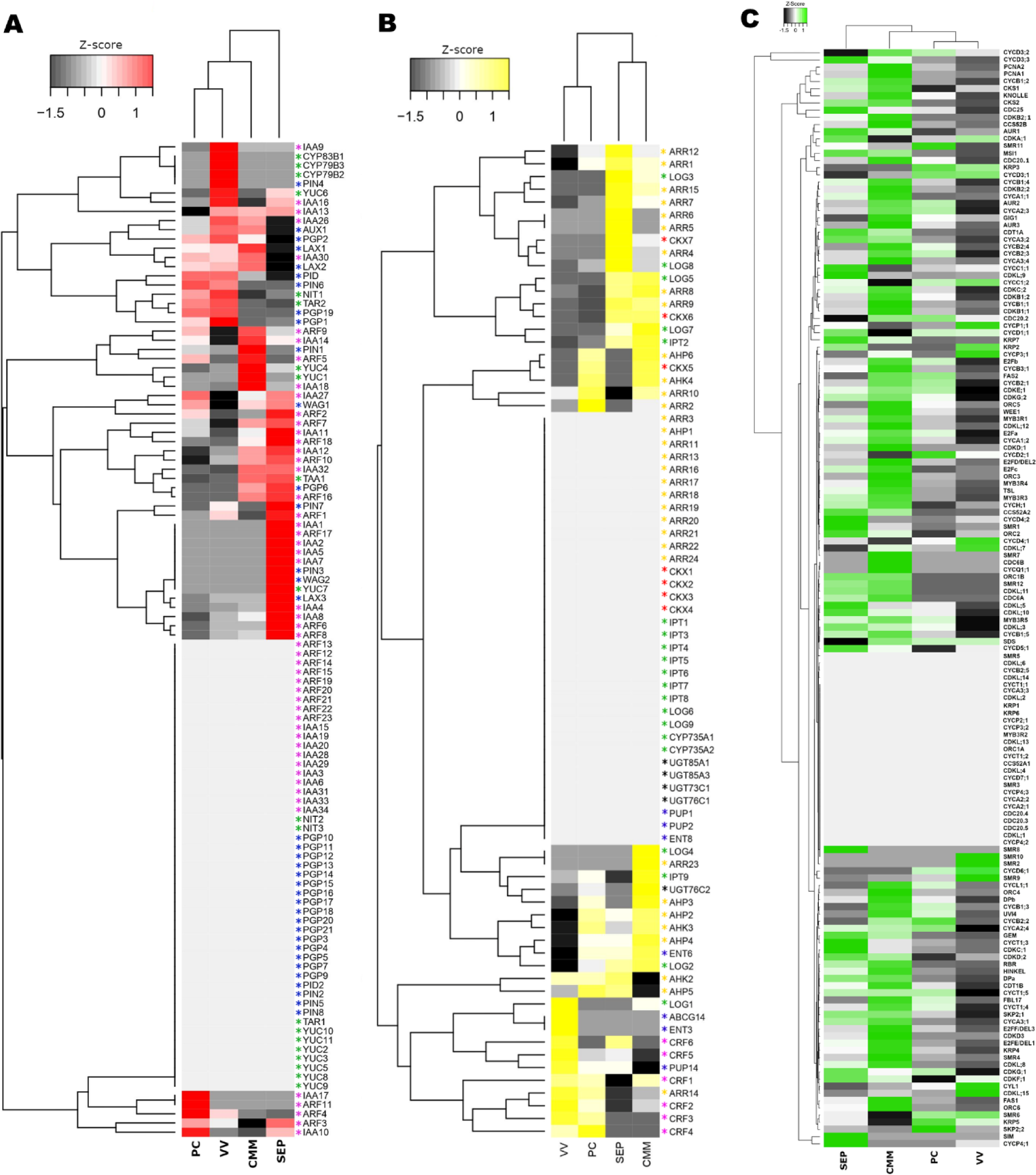
Gene expression dynamics of genes involved in auxin, cytokinin, and cell cycle pathways. **A and B)** Heatmap of the expression profiles of genes involved in auxin and cytokinin homeostasis in the CMM, PC, SEP, and VV tissues. In **A)** colored asterisks: purple – response pathway genes, green – transport pathway genes, blue – biosynthesis pathways genes. In **B)** colored asterisks: purple – response pathway genes, green – biosynthesis pathways genes, blue – transport pathway genes, red – degradation pathways genes, and yellow – signaling pathway genes. **C)** Heatmap of the expression dynamics of cell cycle core genes in the CMM, PC, SEP, and VV tissues.

Previous studies related to cytokinin have reported that expression of the TCS reporter (*TCS::GFP*) is observed in medial tissues such as the CMM, SEP, and transmitting tract (e.g., Marsch-Martínez et al., 2012b; Müller et al., 2017; Reyes-Olalde et al., 2017). As this reporter line has a synthetic promoter containing type-B *ARABIDOPSIS RESPONSE REGULATOR* (*ARR*) binding sites, it is likely that some type-B *ARRs* are expressed in these tissues, and thus should be included in our data set. Indeed, as visualized in Figure 3A, out of the 11 type-B *ARR* genes, *ARR1*, *ARR10*, and *ARR12* are strongly expressed in the medial tissues (CMM and/or SEP). This is consistent with phenotypes observed in the medial domain of the gynoecium in the triple *arr1 arr10 arr12* mutant (Reyes-Olalde et al., 2017). In the latter work, we also reported a connection between the cytokinin signaling pathway and the auxin biosynthesis and transport pathways in the medial domain during gynoecium development. ARR1, together with the transcription factor SPT, activates the auxin biosynthesis pathway, through the activation of *TRYPTOPHAN AMINOTRANSFERASE OF ARABIDOPSIS1* (*TAA1*), a gene that, together with genes of the *YUCCA* (*YUC1*, *YUC4*, and *YUC7*) family, is significantly enriched in the medial domain samples (> 3-fold, considering an FDR < 0.05; CMM and SEP, Figure 3B). Furthermore, ARR1 and SPT regulate the auxin transport pathway by activating the expression of *PIN3*, a gene of the PIN-FORMED (PIN) family of polar auxin transporters that, together with *PIN1* and *PIN7*, are significantly enriched in the medial domain samples (> 3-fold, considering an FDR < 0.05; CMM or SEP). The idea is that the auxin transporters transport the synthesized auxin from the medial domain to the lateral domain, followed by basal-apical auxin transport for gynoecium growth. In the lateral domain (PC and VV), we can see the AGC13-type protein kinase *PINOID* (*PID*) coding gene (> 2-fold enriched, considering an FDR < 0.05), which phosphorylates PIN proteins important for their polar localization. On the other hand, we observe various AUXIN RESPONSE FACTORS (ARFs) in the medial domain, including ARF6 and ARF8, known to be important for pollen transmitting tract tissue formation (Crawford & Yanofsky, 2011).

Taken together, these results indicate strong regulation of auxin and cytokinin pathway genes leading to hormone homeostasis during gynoecium development.

### Dynamics of the expression of core cell cycle genes

Gynoecium development is dynamic and in a short time frame, tissue formation, patterning, and differentiation take place (Deb et al., 2018; Ferrándiz et al., 2010; Herrera-Ubaldo & de Folter, 2022; Moubayidin & Østergaard, 2017; Reyes-Olalde et al., 2013; Roeder & Yanofsky, 2006). During early gynoecium development, in the medial domain, there is the CMM tissue with cells with meristematic activity, which gives rise to the differentiating medial tissues, one of which is the septum (SEP). In the lateral domain, after cell division, cell elongation is a major process. Considering these points, we decided to analyze the transcriptional activity of cell cycle genes during medial and lateral domain tissue development, possibly reflecting their involvement in the progressive development of the gynoecium.

Based on the literature (Gutierrez, 2009; Menges et al., 2005; Shimotohno et al., 2021; Vandepoele et al., 2002), we considered 153 genes part of the core cell cycle (Supplemental Table S16, Figure 3C). Of these 153 genes, more than 110 genes are expressed (≥ 4 TPM), and 26 genes were differentially expressed (considering an FDR < 0.05) in one of the four tissues studied (CMM, PC, SEP, and VV). Each of these 110 cell cycle genes expressed in our data set showed a specific and dynamic pattern in each of the four tissues (Figure 3C).

As mentioned, the CMM cells have meristematic activity and likely a high activity of the cell division cycle; in line with this, many cell cycle genes show a high expression in this tissue (Figure 3C). In comparison with the other tissues, the CMM has the most expressed genes, followed by the septum tissue (SEP). The lateral domain tissues (PC and VV) show less expression of cell cycle genes. The CMM sample shows high expression of all four type-B cyclin-dependent kinases (CDKs): CDKB1;1, CDKB1;2, CDKB2;1 and CDKB2;2, which regulate the G2/M transition (Gutierrez, 2009). Examples of other CDKs higher expressed in the CMM are the CDKC;2, CDKD;1, and CDKD;3 kinases, which, in conjunction with some cyclins, can activate type-A and type-B CDKs.

In addition to these CDKs, CDKL8 and CDKL12, genes belonging to a large group of CDK-like genes, are highly expressed in CMM tissue. On the other hand, the SEP sample is characterized by the expression of five CDKs: one is CDKC;1, and the rest are CDK-like proteins (CDKL;3, CDKL;5, CDKL;9 and CDKL;10). As opposed to this, only CDKL;7 and CDKL;15 are highly expressed in VV tissue during the development of the lateral domain.

In addition to the expression of CDKs, the expression of other genes such as a catalytic CDK subunit (CKS) and cyclins (CYCs) is also necessary, which together these three proteins form an active complex (CDK/CYC). The CKS genes codify for proteins that act as docking factors that mediate the interaction of CDKs with putative substrates and regulatory proteins, and interestingly the two CKS (CKS1 and CKS2) described to date are more expressed in the medial domain (both in CMM and SEP, but mainly in CMM). Related to the CYCs, they are expressed in both the medial and lateral domains; the medial domain is characterized by a large number of highly expressed CYCs. The CMM tissue showed mainly an expression of the CYCA and CYCB groups, which are major CYCs that control the cell cycle during the G2/M transition, such as: CYCA1;1, CYCA1;2, CYCB1;1, CYCB1;2, CYCB1;4, CYCB2;4, CYCB3;1, as well as CYCD3;2, CYCQ1;1 and CYCT1. Meanwhile, in the SEP tissue, the highest expressed CYCs are CYCD members (CYCD3;3, CYCD4;2, CYCD5;1), which also control the cell cycle and whose expressions is regulated by growth-promoting factors such as auxin, cytokinins, and brassinosteroids. In addition, cyclins such as CYCC1;1, CYCP4;1, and CYCT1;3, are also highly expressed in the SEP. However, although in smaller numbers, the CYCs are also expressed in the lateral domain, such as CYCD2;1, the only CYC expressed in the PC tissue, while cyclins CYCD4;1, CYCP1;1, CYCP3;1 and CYL1 are expressed in the VV tissues.

Correct cell cycle progression also involves expression of the E2F transcriptional regulators, which consist of a heterodimer of the related proteins E2F and DP, which when bound to the RETINOBLASTOMA-RELATED (RBR) protein, inhibits its activity. However, when the CDK/CYC complex phosphorylates the RBR protein, RBR dissociates from E2F/DP, and the heterodimer is active and stimulates the transcription of genes needed for cell cycle progression. In Arabidopsis, eight members (DPa, DPb, E2Fa, E2Fb, E2Fc, E2F/DEL1, E2F/DEL2 and E2F/DEL3) make up the E2F family, and interestingly, all of them are higher expressed during medial domain development compared to the lateral domain. In the medial domain, we observed that their expression is higher early in development (CMM) compared to a later development stage (SEP). Interestingly, RBR expression also shows this tendency, suggesting an interaction between the E2F transcription factors, RBR, and CDK/CYC complex during gynoecium development, mainly in the CMM tissue, to control meristem activity.

Lastly, the molecular machinery that controls cell proliferation is under strong regulatory control. For example, the activity of CDK/CYC complexes is repressed by CDK inhibitors (CKIs), which are divided into two main groups, one is the interactor/inhibitor of the cyclin-dependent kinase/KIP-related protein (ICK/KRP) family, which comprises 7 members (KRP1-KRP7). Of these, KRP4, KRP7, and KRP2 are highly expressed in the medial domain, while KRP5, KRP3, and KRP2 are highly expressed in the lateral domain (PC and VV). The other group consists of plant-specific Siamese (SIM) and Siamese-related (SMR) proteins. During the development of the medial domain, SMR12 is expressed both in CMM and SEP tissues, but only SMR4 and SMR7 are highly expressed in CMM. SIM, the main regulator of this group, is highly expressed in the SEP along with SMR1 and SMR8. On the other hand, the lateral domain is characterized by the expression of SMR6 in both PC and VV, and the SMR11 gene is more expressed in PC tissue, while the SMR2, SMR9, and SMR10 genes are highly expressed in VV tissues.

In summary, the cell division cycle is clearly highly dynamic and likely to be very important for gynoecium development.

### Identification and analysis of differentially expressed genes

To identify differentially expressed genes (DEGs), one-versus-all tissue comparisons were made; CMM was compared against SEP+PC+VV, and with the same logic, each tissue was compared to the other three tissues using the R package edgeR (Robinson et al., 2009). A gene was considered a DEG, if it had a fold change >1 and a False Discovery Rate (FDR) < 0.05. As a result of these analyses, for the medial domain tissues, a total of 3,648 and 2,416 DEGs were statistically significant in the CMM and SEP, respectively, while 1,569 and 4,119 DEGs were statistically significant in the lateral domain PC and VV, respectively (Figure 4A, Supplemental Tables S17-S20).

**Figure 4.**
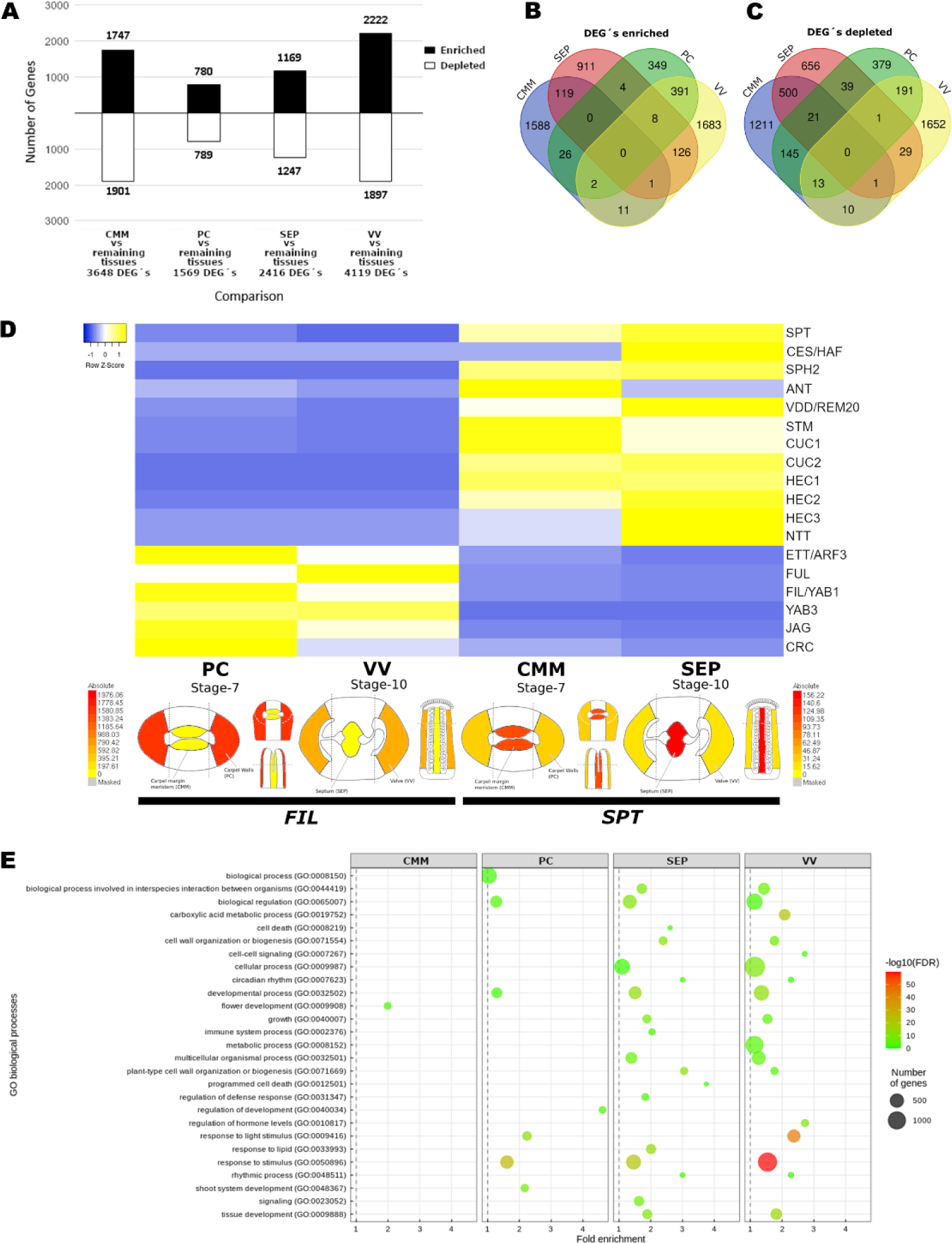
Differential gene expression analysis. **A)** Number of DEGs of each tissue. **B and C)** Venn diagrams of enriched and depleted DEGs (fold change > 1 and FDR < 0.05), showing the unique and common DEGs between the CMM, PC, SEP, and VV tissues. **D)** Heatmap showing the expression profiles of transcription factors (which are DEGs in of the tissues) previously reported to regulate the development of the medial (CMM and SEP) and lateral domain (PC and VV) tissues of the gynoecium. Below the heatmap, eFP Browser views of expression profiles for *FILAMENTOUS FLOWER* (*FIL*), a DEG of the lateral domain, and *SPATULA* (*SPT*), a DEG of the medial domain; these views exemplify the visualization of our data available in the Arabidopsis eFP Browser (Winter et al. 2007). **E)** GO analysis of DEGs of each tissue, applying the methodology of Bonnot et al. 2019.

Next, we wanted to know how many of the DEGs were genes coding for transcription factors. For this, we graphed all unique transcription factors of the DEGs of each tissue together in a circle plot and then highlighted each unique transcription factor present in a specific tissue (Supplemental Figure 3). In each circle plot, the percentage is given of how many of the total number of Arabidopsis transcription factors are part of the DEGs in each tissue: 8.2% in CMM, 6.2% in SEP, 2.9% in PC, and 8.7% in VV tissue (Supplemental Figure 3 and Supplemental Table S21).

In an overlap analysis of the DEGs of the four tissues, we were able to identify shared DEGs between tissues that were enriched (increased) and depleted (decreased) in expression (Figures 4B and 4C, respectively). Interestingly, the most overlap is found between the tissues of the same domain of the gynoecium, although they are in different developmental stages, i.e., between CMM and SEP (medial domain), and between PC and VV (lateral domain). Furthermore, we also identified unique DEGs for each tissue that were enriched or depleted.

To obtain insight into the specificity of LAM dissected material and the reliability of the identified DEGs, we searched the list of enriched DEGs of each tissue (Supplemental Tables S17-S20) for the presence of known transcription factors involved in the development of that particular tissue or domain (Figure 4D, Supplemental Table S22). Some “marker” genes found for medial domain development were: *SPT*, *CES/HAF*, *SHATTERPROOF2* (*SHP2*), *ANT*, *VERDANDI* (*VDD/REM20*), *STM*, *CUC1* and *CUC2*, *HEC1, HEC2* and *HEC3* and *NTT*. For lateral domain development, we found “marker” genes such as *ETT*, *FUL*, *FIL*, *YABBY 3* (*YAB3*), *JAG*, and *CRC* (Figure 4D, Supplemental Table S22).

To analyze the biological processes involved in medial and lateral domain development, a Gene Ontology (GO) enrichment analysis was performed for the enriched DEGs of each tissue, for which we used the Bonnot and collaborators (2019) method (Bonnot et al., 2019). After using the PANTHER and the REVIGO platforms, we recovered and plotted the significantly enriched functional and process terms (Figure 4E, Supplemental Table S23), such as flower development (GO:0009908), the only GO recovered for the CMM, while for SEP we found the categories cell death (GO:0008219), programmed cell death (GO:0012501), response to lipid (GO:0033993), and signaling (GO:0023052). In contrast, developmental process (GO:0032502), regulation of development (GO:0040034), and response to light stimulus (GO:0009416) were enriched terms for PC, while for VV, we recovered cell-cell signaling (GO:0007267), metabolic process (GO:0008152), regulation of hormone levels (GO:0010817), and response to light stimulus (GO:0009416). These GO results suggest that the enriched DEGs consist of genes that change over developmental time and are distinct in each domain of the gynoecium.

### Promoter analysis of DEGs

To obtain insights into what transcription factors could regulate the identified DEGs in each tissue, an *in silico* analysis was performed to identify statistically significant DNA-binding motifs enriched in the upregulated DEGs of the CMM, PC, SEP, and VV tissues (we used 500 bp upstream of the ATG of the promoter regions of these genes). In general, DNA-binding motifs of the TCP, WRKY, bHLH, MYB, and AHL transcription factor families were enriched in all tissues (Figure 5). On the other hand, the DNA-binding motifs with the highest percentage of occurrence in the CMM tissue were the YABBY family of transcription factors (YAB1 and YAB5), WRKY (WRKY12,18,38, and 45), AP2 (TOE1), and the R2R3-MYB transcription family (MYB111, 56, 55, and 59) (Figure 5A), and, interestingly, the DNA-binding motifs of TOE1 and WRKY18 are only enriched in the CMM tissue (Figure 5A, B). In the PC tissue, members of the AT-hook (AHL12 and AHL25), R2R3-MYB (MYB55), bHLH (MYC3), and the YABBY (YAB5) family were the most enriched DNA-binding motifs. The SEP tissue is enriched in DNA-binding motifs such as family members of AHL (AHL20 and AHL25), HD-Zip I (ATHB12 and ATHB51), R2R3-MYB (MYB45, MYB52, MYB111, and MYB59), as well as the superfamily B3, which includes the DNA-binding motif of REM1, which is enriched only in septum tissue (Figure 5A, C). In the VV tissue, we found enrichment of the DNA binding motifs of the families AT-hook (AHL20, AHL25), ARR type-B (ARR14), ARF (ARF3/ETT), as well as MYB55, PIF5, WOX13, REM1_2 (splicing version of REM1), SPL1 and ZAT2 transcription factors (Figures 5A, D). Lastly, three binding motifs, Dof5.7, ARR11, and RRTF1, were enriched in CMM and SEP tissue (Figure 5E).

**Figure 5.**
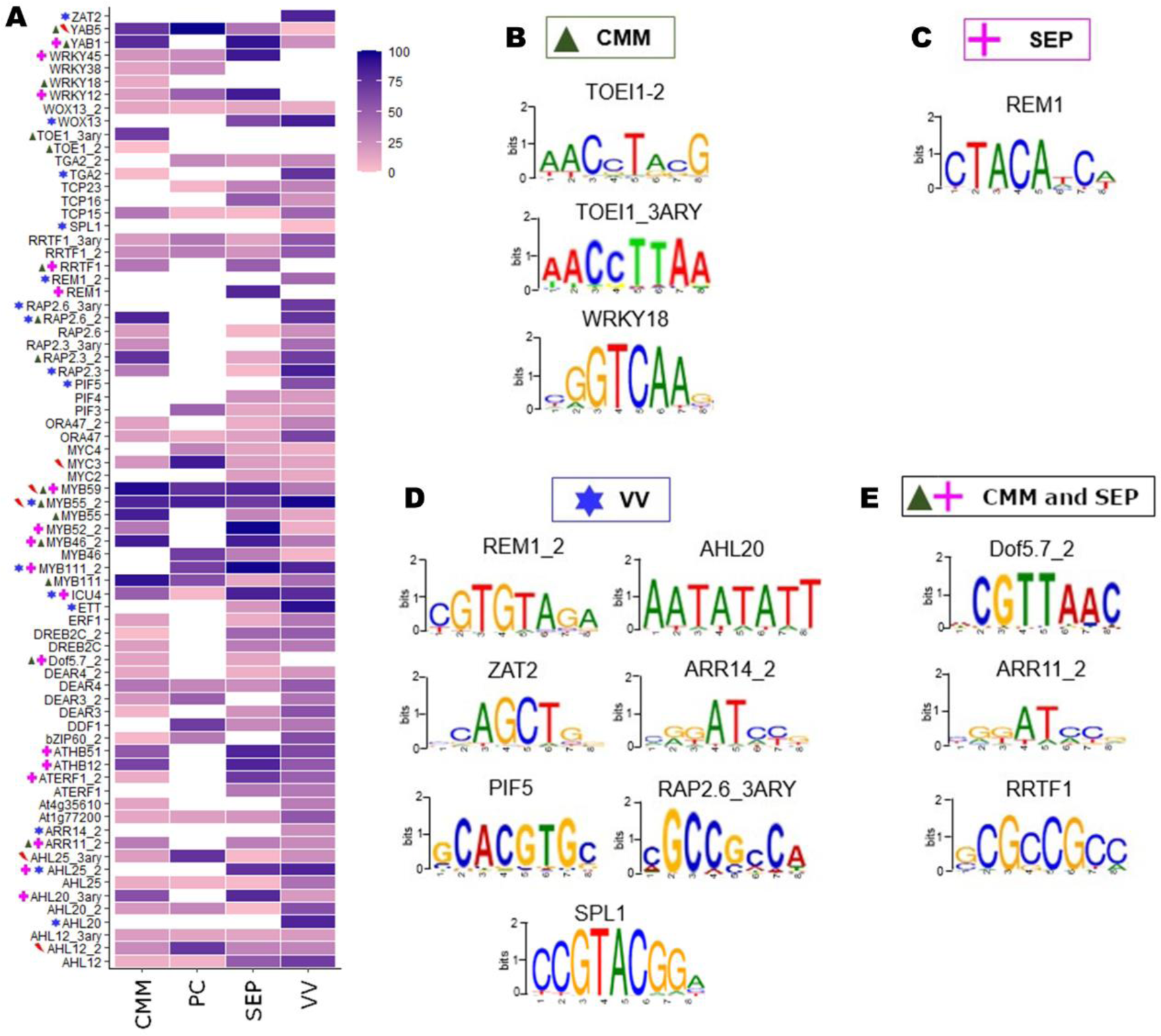
Promoter DNA-binding motif enrichment analysis of DEGs. **A)** DNA-binding motif enrichment analysis in the promoter of DEGs of each tissue. Scale bar indicates the percent of enrichment of the motif in the total of upregulated DEGs. Symbols next to the heatmap indicate the DNA-binding motif with ≥70% occurrence in the promoter region of upregulated DEGs or unique DNA-binding motifs found in CMM (green triangle), PC (red lightning), SEP (pink cross), or VV (blue star). **B to E)** Unique DNA-binding motifs found in promoters of DEGs of CMM, SEP, and VV. In PC tissue, no unique DNA-binding motifs were detected.

### A functional analysis to identify new regulators of gynoecium development

The results of the generation of an expression atlas and the presented transcriptomic analyses have allowed us to elucidate the transcriptional dynamics that regulate the development of some tissues that make up the medial and lateral domains of the gynoecium. Besides the presence of known genes and transcription factors involved in gynoecium development, we also observed many genes present in the lists of DEGs that have not yet been studied in gynoecium development. We decided to study several of these genes classified as DEGs of the medial domain, and we report here the results of mutants of 5 genes of the CMM and 2 genes of the SEP tissue (Figures 6A and 7A, G). For the CMM tissue these are: *AT1G51460* (*ABCG13 - ATP-BINDING CASSETTE G13, FOP2 - FOLDED PETALS 2*), a gene that encodes a member of the ATP-binding cassette (ABC) transporter family protein (Panikashvili et al., 2011; Takeda et al., 2014); *AT1G76540* (*CDKB2;1* - *CYCLIN-DEPENDENT KINASE B2;1*), a gene that encodes a cyclin-dependent protein kinase involved in the regulation of the cell cycle (Boudolf et al., 2001; Vandepoele et al., 2002); *AT3G19184* (*REM1* - *REPRODUCTIVE MERISTEM 1*), a gene encoding a transcription factor of the REM family involved in meristematic tissues development (Franco-Zorrilla et al., 2002); *AT2G28610* (*WOX3 - WUSCHEL RELATED HOMEOBOX 3*, *PRESSED FLOWER – PRS*) and, *AT5G17810* (*WOX12*), both members of the *Wuschel-related homeobox* gene family with functions from meristem maintenance to patterning (Liu et al., 2014; Matsumoto & Okada, 2001). For the SEP tissue, we also chose one *WOX* gene, *AT3G18010* (*WOX1*), and the gene *AT5G46700* (*TORNADO 2* [*TRN2*] / *TETRASPANIN 1* [*TET1*]); the latter being a member of a membrane protein family that has functions in cell adhesion, fusion, polar growth, membrane trafficking, signaling, motility, and morphogenesis (Chiu et al., 2007).

**Figure 6.**
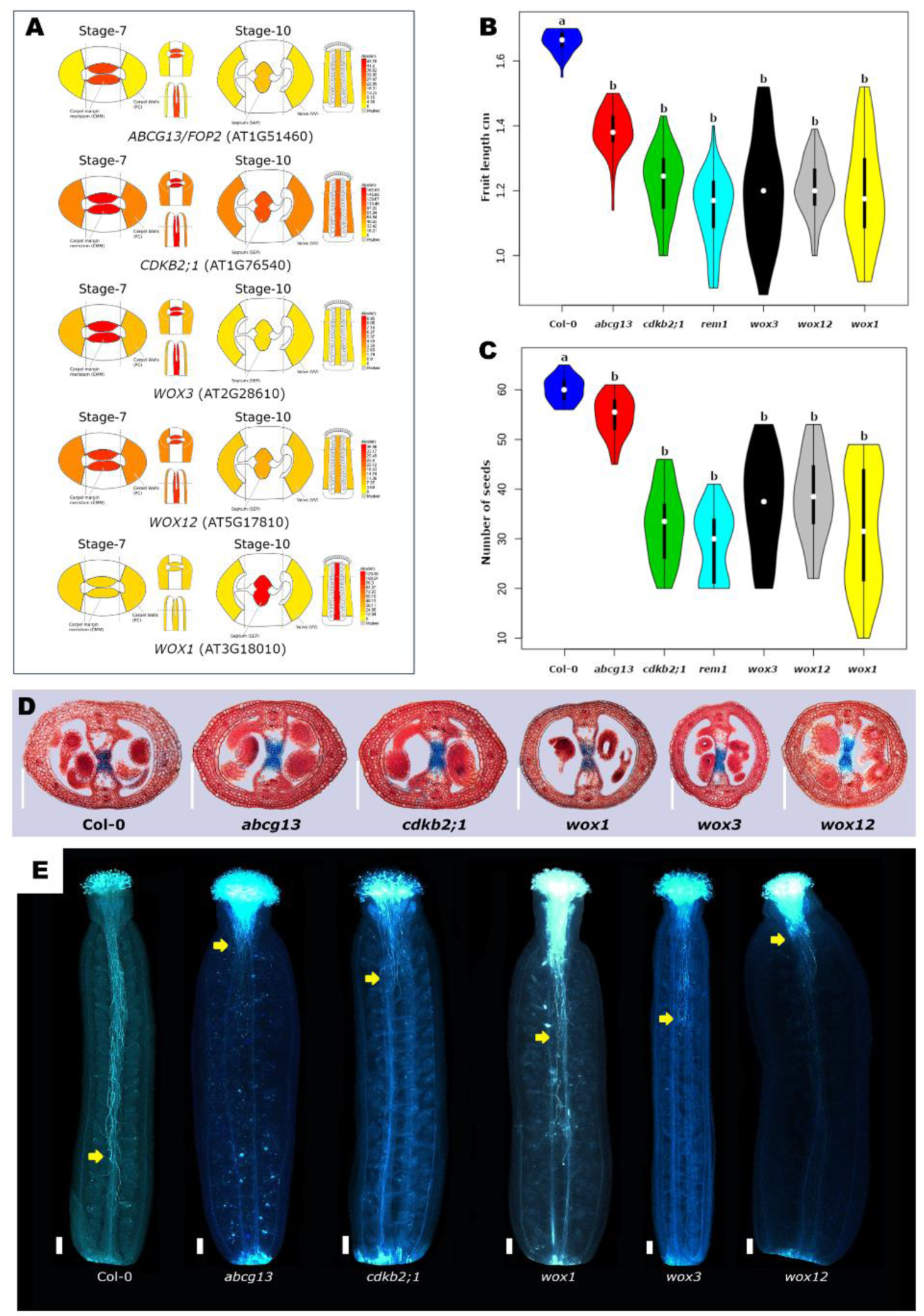
Functional analyses of candidate genes of medial domain development. **A)** Arabidopsis eFP Browser view of the gene expression profile of *ABCG13*/*FOP2*, *CDKB2;1*, *WOX3*, *WOX12*, and *WOX1*. **B and C)** Quantitative analyses of fruit length and seed number, respectively. Data is represented with violin plots, showing the median (white dot), the interquartile range (the black bar in the centre of the violin), the lower/upper adjacent values (the black lines stretched from the bar), and the frequency values (different colours for each mutant). Different letters above the plots indicate a statistically significant difference, based on an analysis of variance (ANOVA) followed by Tukeýs honest significance (TukeyHSD) test. *n*=50 fruits, p-value <0.0001. **D)** Gynoecia cross-sections stained with alcian blue and neutral red staining of wild type (Col-0), *abcg13*, *cdkb2;1*, *wox1*, *wox3*, and *wox12* mutants at stage 12. Scale bars are 10 µm. **E)** Pollen tubes stained with aniline blue of wild type, *abcg13*, *cdkb2;1*, *wox1*, *wox3*, and *wox12* mutants. Scale bars are 10 µm. The yellow arrows indicate the location up to where the growth of most of the pollen tubes was observed.

**Figure 7.**
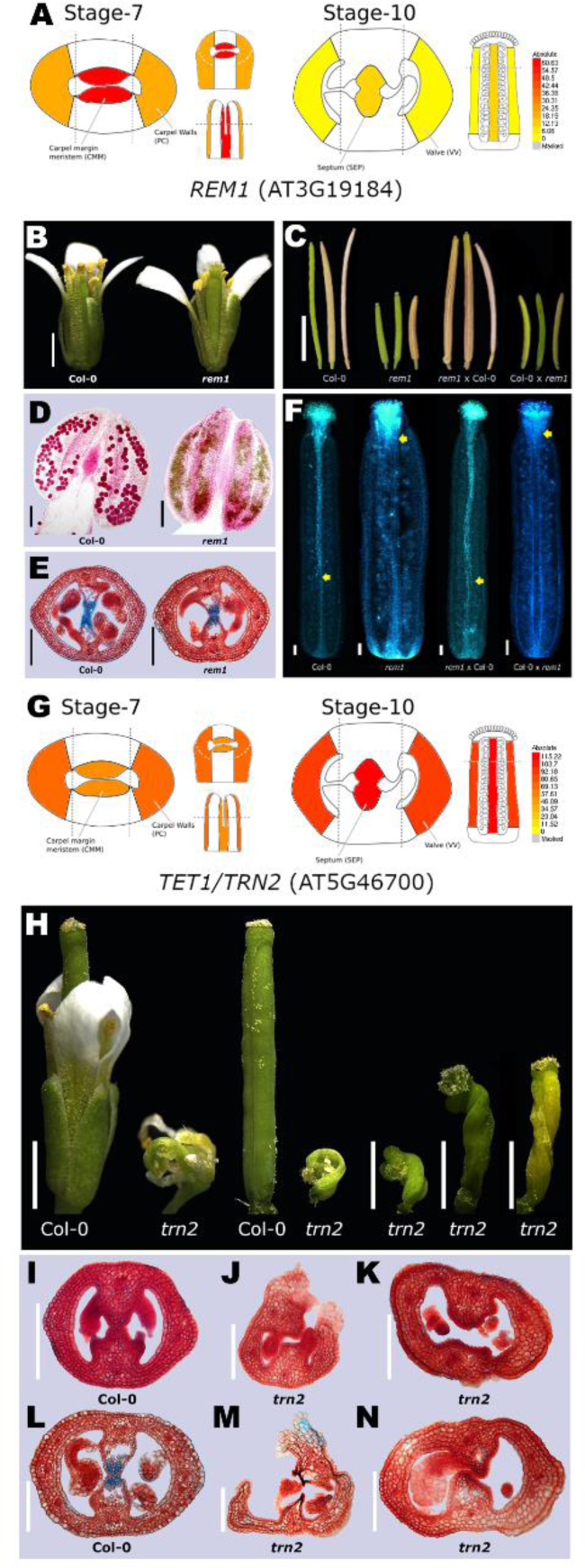
Functional gene analyses of *REM1* and *TET1*. **A)** Arabidopsis eFP Browser view of the gene expression profile of *REM1*. **B)** Wild type and *rem1* flowers. Scale bars are 0.5 cm. **C)** Fruits after reciprocal crosses; Col-0 and *rem1* self-pollination, *rem1* x Col-0, Col-0 x *rem1*. Crosses are indicated female x male. Scale bar is 0.5 cm. **D)** Pollen viability analysis by Peterson’s staining, *rem1* showed aborted pollen, grains colored green. Scale bar is 10 µm. **E)** Alcian blue and neutral red staining of gynoecia cross-sections of wild type and *rem1*, no obvious defects were observed. Scale bars are 10 µm. **F)** Pollen tubes stained with aniline blue, wild type and *rem1* were self-pollination, cross *rem1* x Col-0, and Col-0 x *rem1*. Scale bars are 10 µm. The yellow arrows indicate the location up to where the growth of most of the pollen tubes was observed. **G)** Arabidopsis eFP Browser view of gene expression profile of *TET1*/*TRN2*. **H)** Wild type and *trn2* flower and fruits. Scale bars are 0.5 cm. **I to N)** Alcian blue and neutral red staining of stage 10 gynoecia cross-sections of wild type and *trn2* (**I-K**); unfused septum visible in *trn2* (**J and K**). Stage 12 gynoecia cross-sections of wild type and *trn2* (**L to N**), no transmitting tract is visible in *trn2* (**M and N**). Scale bars are 10 µm.

T-DNA insertion lines for each gene were analyzed (Supplemental Table S24), and all mutants showed defects in reproductive development. In general, all mutants developed shorter fruits with a decreased number of seeds compared to the wild type (Figures 6B-C, Supplemental Figure 4A). In addition, in the *tet1*/*trn2* mutant plants, we observed severely affected flowers with twisted gynoecia and twisted fruits, as reported previously (Figure 7H; Chiu et al., 2007).

To obtain more insights into the role of each gene, we analyzed various aspects of male and female reproductive development in the mutants. First, pollen viability was analyzed with a staining method (Peterson et al., 2010), and only in the *rem1* mutant pollen abortion was observed (Figures 7B-D). Reciprocal crosses of *rem1* with wild type confirmed a pollen defect (Figures 7C, F). The pollen of the other mutants was all viable as determined by the staining of the pollen (data not shown). On the female side, we analyzed gynoecium patterning and development, with a focus on septum and transmitting tract formation in gynoecia cross-sections from each mutant (Figures 6D and 7E, I-N). Only in the *tet1*/*trn2* mutant was a lack of septum fusion and transmitting tract formation observed (Figure 7I-N), which is in line with *TET1*/*TRN2* being a DEG selected from SEP tissue. In the gynoecia of the other mutants, no obvious defects were observed compared to the wild type (Supplemental Figure 4B). Finally, using aniline blue staining we analyzed pollen tube growth (using pollen from the same mutant, as well as pollen from the wild type) in the gynoecium of the mutants (except for *tet1*/*trn2*, due to the severely affected flower). Surprisingly, for the mutants *abcg13*, *cdkb2;1*, *rem1*, *wox3*, and *wox12* pollen tube growth was mainly observed in the apical part of the ovary (Figures 6E, 7F), while in the *wox1* mutant there was a reduced number of pollen tubes in the ovary compared to the wild type (Figure 6E). In addition, when using wild type pollen for pollination on mutant gynoecia of *abcg13*, *cdkb2;1*, *wox1*, *wox3*, and *wox12*, no pollen tube growth was observed (Supplemental Figure 4C), suggesting a problem on the gynoecium side. As expected, using wild type pollen for pollination on mutant gynoecia of *rem1*, pollen tube growth was observed (Figure 7F), suggesting that in this mutant the gynoecium seems to be unaffected.

Altogether, these observations can explain the decreased seed production and, as a consequence, the shorter fruits observed in the mutants.

## Discussion

Multicellular organisms are composed of diverse cell types that are defined by specific transcriptomic, proteomic and metabolomic profiles, and defining them has become a priority in order to understand the molecular processes that drive their development. An essential step on this path is the generation of atlases of gene expression in specific cells of an organism.

In this work, we used Laser-Assisted Microdissection (LAM) microscopy coupled to RNA-Seq to generate a high-resolution gene atlas of the medial and lateral domains of the gynoecium of *Arabidopsis thaliana*. We generated and analyzed gene expression profiles for the medial and lateral domain tissues of the gynoecium at two developmental stages. We sampled at floral stage 7 the carpel margin meristem (CMM; medial tissue) and the presumptive carpel walls (PC; lateral tissue) and at floral stage 10 the septum (SEP; medial tissue) and the valves (VV; lateral tissue).

We found that around half of all Arabidopsis genes are expressed in each tissue, of which around 10% to 30% are differentially expressed between tissues. In general, expression levels are highly dynamic during gynoecium development. Furthermore, we also detected a small number of genes with tissue-specific expression, which can be used in future experimental designs, as markers of the development of these tissues. Notably, tissue-specific gene expression levels were around 1/3 of non-tissue-specific genes. We detected almost all known genes coding for transcription factors and other proteins important for gynoecium development (Reyes-Olalde et al., 2013). The spatial distribution of the expression of the known genes was nicely recovered in our study. All the data and its visualization are available to the community in the Arabidopsis eFP browser (Winter et al., 2007). Furthermore, for additional confidence in our data, we compared the log fold change (logFC) values of the DEGs of each tissue with the published list of DEGs obtained from cells with the medial domain expressed *SHATTERPROOF2* promoter fused to YFP using a FACS-based system (Villarino et al., 2016). Based on a Spearman rank correlation analysis, the CMM and SEP medial tissues (in this study) clustered with the medial expressed genes of the SHP2-YFP positive cells (Supplemental Figure 5; Villarino et al., 2016).

Related to two important hormones, cytokinin, and auxin, homeostasis seems to be a complex process where they act cooperatively to regulate gynoecium development. Around 56% of the genes that have been reported to be involved in biosynthesis, transport, signaling, and response pathways for both hormones are differentially expressed in the four tissues analyzed, including the genes previously reported by Reyes-Olalde et al., (2017) to be involved during the development of the gynoecium. Furthermore, more genes that are part of these hormonal pathways show interesting expression patterns that have not yet been studied related to gynoecium development, as well as genes related to other hormonal pathways.

On the other hand, in the analysis of cell cycle genes, although a large number of genes are expressed in all the tissues analyzed, the development of the medial domain is characterized by the expression of the two subgroups of B-type cyclin-dependent kinases, CDKB1 and CDKB2 (CDKB1;1, CDKB1;2, CDKB2;1, and CDKB2;2), which are expressed during S to M phase and G2 to M phase, respectively, in cell cycle progression (Inzé & De Veylder, 2006). As they are known to regulate the stem cell niche organization in the shoot apical meristem (Andersen et al., 2008), this may suggest that the medial domain, which contains the CMM, is in constant cell cycle progression.

The *in silico* analysis of DNA-binding motifs in the promoters of DEGs in each tissue showed enrichment of distinct members of transcription factor families. Several members of the TCP and bHLH transcription factor families were found to be enriched in tissues from CMM, PC, SEP, and VV, which is interesting and consistent, as members of both families have been reported to be expressed and to be involved during the medial and lateral domain development of the gynoecium (Alvarez & Smyth, 1999; Cubas et al., 1999; Di Marzo et al., 2020; Foreman et al., 2011; Girin et al., 2011; Groszmann et al., 2008; Heisler et al., 2001; Liljegren et al., 2004; Lucero et al., 2015; Nahar et al., 2012; Rajani & Sundaresan, 2001; Reyes-Olalde et al., 2017; Reymond et al., 2012). On the other hand, the DNA-binding motif of REM1 was observed to be enriched during medial domain development, specifically in SEP tissue; interestingly, the gene *REM1* is differentially expressed in the medial domain. The REM family of transcription factors is known to be expressed in reproductive meristems (Franco-Zorrilla et al., 2002; Mantegazza et al., 2014; Villarino et al., 2016). Also, DNA-binding motifs of transcription factors involved in cytokinin signaling (ARR14 and ARR11) were enriched in the medial tissues, which are known to be involved during the development of these tissues (Herrera-Ubaldo et al., 2023; Marsch-Martínez & de Folter, 2016). In the lateral tissues, DNA-binding motifs of members of the auxin family (ARF4) were enriched, supporting the function that has been attributed to this transcription factor, and its close homolog ARF3 (ETTIN), in gynoecium patterning (Nemhauser et al., 2000; Pekker et al., 2005).

In the last decades, numerous transcription factors have been identified and connected to be important for gynoecium development (Herrera-Ubaldo & de Folter, 2022), and novel genes and connections are still being discovered (Herrera-Ubaldo et al., 2023; Ramos-Pulido & de Folter, 2023). In this work, based on transcriptome data, new gene functions in gynoecium development were also discovered. We analyzed mutants for seven genes and found that all are involved in reproductive development, ranging from mild to severe phenotypes.

In summary, this study provides a tissue-specific gene expression analysis of the medial and lateral domains of the Arabidopsis gynoecium, and together with the results of the functional gene analyses, shows itself to be a useful resource for the study of reproductive development.

## Material and Methods

### Plant material and growth conditions

*Arabidopsis thaliana* plants wildtype Col-0 and T-DNA insertion lines were germinated in soil (3:1:1, peat moss:perlite:vermiculite) in a growth chamber under long-day conditions (16 hours light - 8 hours dark) for 2 weeks and transferred to standard greenhouse conditions (22–27°C, natural light). All mutant lines (*abcg13* or *fop2-2*, SALK_046735C; *cdkb2;1*, CS874085; *rem1*, CS869296; *tet1*, SALK_127323C; *wox1*, SALK_148070C; *wox3*, CS863894; *wox12*, SALK_087882C) were obtained from the Arabidopsis Biological Resource Center (ABRC).

### Tissue embedding for laser assisted microdissection (LAM)

The inflorescence embedding protocol is based on the protocol described by Chávez Montes et al. (2016). In summary, open flowers were removed from each main inflorescence. These main inflorescences were harvested in ice-cold fixation solution of ethanol 100%:acetic acid, (9:1-v/v ratio), infiltrated twice under vacuum conditions for 15 min and stored overnight in new fixation solution at 4°C. The fixation solution was removed, and the inflorescences were dehydrated with a graded ethanol series (ethanol 70, 80, 90, 100, 100%) at 4°C, shaking (Mini-rotator, Glas-Col, USA) at 20 rpm, with each step taking 1 h, and then stored overnight in 100% ethanol at 4°C without shaking. The next day, the ethanol was slowly replaced by histoclear using the following series: ethanol:histoclear (v:v) 3:1, 1:1, 1:3, 100% histoclear, 100% histoclear, with each step taking 1 h, at room temperature with mild and occasional shaking. Finally, the inflorescences were stored overnight in pure histoclear with paraplast beads (1:1, v/v) at room temperature. The next day, inflorescences were moved to 58°C for 10 ∼ 15 min, then the mixture of histoclear-paraffin beads was gradually removed and replaced with liquid paraffin (first 25% of the total volume of the mixture of histoclear-paraffin was decanted and replaced by pure liquid paraffin, this was repeated at 50%, 75%, 100%, 100% of total volume), the liquid paraffin was changed every 1 ∼ 1.5 h at 58°C. Finally, the inflorescences were stored overnight in 100% liquid paraffin at 58°C. Finally, the blocks were assembled with liquid paraffin and 3∼4 flowers embedded per block. The blocks were assembled and allowed to solidify on ice and stored at 4°C.

### Laser assisted microdissection

The embedded tissue in paraffin blocks were transverse sectioned 12 µm-thick with a Leica RM2035 microtome (Leica, Germany) and sections were fixed on slides with pure methanol at 45°C for ∼45 min. The slides with sections were dewaxed for 15 min in histoclear. After that, the slides with sections were dried at room temperature for 15 min, and then the tissues of interest were immediately microdissected with a Zeiss PALM MicroBeam IV laser microdissection microscope (Zeiss, Germany).

### RNA extraction, cDNA synthesis and library preparation

For each sample, microdissected material of around 80 gynoecia (sections of around 24 paraffin blocks with 3 or 4 embedded inflorescences) was pooled in a single tube and RNA extraction was performed with Direct-zol RNA MicroPrep kit (Zymo Research). The integrity and concentration of extracted RNA of each sample was measured using a NanoDrop 2000 (ThermoFisher Scientific). For each tissue, three biological replicates of RNA were collected, generated from independent plant material for each sample. The sequencing libraries were prepared from 10 ng total RNA input per sample, for cDNA synthesis the SMART-Seq v4 Ultra Low Input RNA Kit for Sequencing (Takara Bio) was used, as specified by the manufacturer’s instructions. All synthesized cDNA was fragmented by sonication using the Covaris S2 Focus Ultrasonicator (Covaris) to generate cDNA fragments of ∼100 bp.

For library preparations, the NEBNext Ultra II DNA Library Prep Kit for Illumina (New England Biolabs) and the NEBNext Multiplex Oligos for Illumina (Index Primers Set 1; New England Biolabs) were used as specified by the manufactureŕs instructions. The input material for each library was all previously synthesized and fragmented cDNA. Sequencing was conducted at Novogene (California, USA) using the 150 paired-end Illumina HiSeq2000 sequencing platforms.

### Mapping reads to the reference transcriptome

The quality of raw sequencing reads was analyzed by FASTQC v0.11.5 (Andrews, 2010) and overrepresented sequences, low-quality reads and adapters were removed using trimmomatic v0.39 (Bolger et al., 2014) and cutadapt v2.8 (Martin, 2011). The sequencing reads filtered (Phred Quality Score >30) were mapped to the *Arabidopsis thaliana* reference transcriptome (Araport11, https://www.arabidopsis.org/download/index-auto.jsp?dir=%2Fdownload_files%2FSequences%2FAraport11_blastsets) using Kallisto v0.46.1 (Bray et al., 2016). Read counts and transcript per million reads (TPMs) were generated using the R package tximport v1.0.3 and the lengthScaledTPM method (Soneson et al., 2015).

### Gene expression and differential gene expression

Gene expression and differential gene expression (DEG) analyses were carried out using R packages tximport v1.12.3 (Soneson et al., 2015) and edgeR v3.26.8 (Robinson et al., 2009). First, to know which genes were expressed in each tissue, the TPM (Transcripts Per Kilobase Million) values were used to represent the expression level of each gene per tissue. To consider that a gene had expression, it had to have an expression > 0TPM in each of the three biological replicates of each tissue analyzed, in addition the average expression of the three biological replicates had to be > 4TPM. The gene expression values were used to make a heatmap with hierarchical clustering of genes expressed in the four tissues, for this goal the k-means and heatmap.2 functions of R were used.

The remaining heatmaps were built using the package ggplots and heatmap.2 function of R.

For DEG analysis we used normalized data and the comparisons that were made were one tissue type vs all remaining tissues. For this analysis, lowly expressed transcripts were filtered based on analyzing the mean-variance trend, and transcripts with more than 1 counts per million reads were retained. Reads were normalized using the TMM method and a linear model of the negative binomial was made to characterize count data and allow for multi-factor design, furthermore, the empirical Bayes shrinkage was used to estimate the dispersion parameter and the likelihood ratio test to obtain the p-value. The Benjamini-Hochberg procedure was used to adjust p-values to control the False Discovery Rate (FDR). To consider a differential expressed gene, we apply the cutoff value of a log2 fold change (logFC) > 1 and a FDR < 0.05.

### Arabidopsis eFP Browser: visualization of all expression data

In order to develop an interactive way to study the genes, the raw counts were normalized (using the MEVlab platform) to FPKM (Fragments Per Kilobase Million) and these new values are available in the Arabidopsis eFP browser platform (Winter et al., 2007).

### Gene set enrichment analysis

The Gene Ontology term analysis was performed using the method of Bonnot et al. (2019). In brief, The Arabidopsis Information Resource (TAIR) for GO Term Enrichment for Plants interface (https://www.arabidopsis.org/tools/go_term_enrichment.jsp), which is fed by Panther classification system was used to obtain GO terms, where we choose a Fisheŕs Exact test and a correlation of Calculated False Discovery Rate (FDR). Based on the above results, we identified the most representative GO terms using REVIGO web server (Supek et al., 2011) (http://revigo.irb.hr/), which removes redundant terms and we kept the terms with a dispensability <0.05 to plot. From this list of selected GO terms, we looked for matches in the PANTHER result list and retrieved corresponding GO biological process, the number of genes, fold enrichment and FDR. From this last list, we choose only the over-represented GO terms (GO terms with fold enrichment > 1), which were plotted in RStudio using the ggplot2 package.

### Promoter analysis of DEGs

For promoter analysis, we used MEME suite 5.5.0 (https://meme-suite.org). We extracted from TAIR (https://www.arabidopsis.org) the 500 bp nucleotide sequence upstream of the ATG of each induced tissue-specific DEG (CMM, SEP, PC, VV). Then we used the Simple Enrichment Analysis of motifs (SEA) (Bailey & Grant, 2021) with the motif database of (Franco-Zorrilla et al., 2014) with an E-value ≤10 score threshold and adjusted *p*-value.

### Selection of mutants

T-DNA SALK insertion mutants were obtained from the Arabidopsis Biological Resource Center. To obtain homozygous mutants for every gene, genotyping was performed for the *rem1*, *wox1*, and *wox12* mutants by PCR using the primers mentioned in Supplemental Table S24. The *abcg13*, *tet1/trn2* and *wox3* have been described previously (Chiu et al., 2007; Matsumoto & Okada, 2001; Panikashvili et al., 2011; Takeda et al., 2014).

### Statistical analysis

For the statistical analysis of fruit length and number of seeds, we collected 10 fruits from 5 homozygous plants of each mutant (*n*=50), the fruits were collected on the main stem from fruit number 6 onward of each plant. The fruit length measurements were performed with ImageJ, and from the same fruits the number of seeds were counted. The data were processed by an analysis of variance (ANOVA) followed by Tukey HSD (Honestly Significant Difference) test, using Rstudio version 3.6.3.

### Histology and microscopy analyses

Pollen vitality was analyzed using the Peterson’s stain (Peterson et al., 2010). Flowers between stages 12 and 13 were collected (when pollen was mature but anthers non-dehiscent) and fixed in Carnoy’s fixative (6 alcohol:3 chloroform:1 acetic acid) for a minimum of 2 h. After individual anthers were dissected and placed on a microscope slide, 2-4 drops of the staining solution were added. Then the slide was slowly heated the slide over an alcohol burner in a fume hood until the staining solution was near boiling (∼30 sec). Finally, a coverslip was placed over the sample. Pictures were taken with a DM6000B microscope (Leica).

For transverse gynoecia section analysis, we followed the method previously described (Herrera-Ubaldo et al., 2019). In summary, inflorescences of wildtype and all mutants were collected and fixed in FAE solution (3.7% formaldehyde, 5% glacial acetic acid and 50% ethanol) with vacuum (20 min, 4°C) and incubated for 2 h at room temperature. The material was rinsed with 70% ethanol and incubated overnight at 4°C in 70% ethanol, followed by dehydration in a series of ethanol dilutions (70%, 85%, 95% and 100% ethanol) for 1 h each. Inflorescences were embedded in Technovit 7100 (Heraeus Kulzer) according to the manufacturer’s instructions. Sections (12-15 μm thick) were obtained on a rotary microtome (Leica). To observe the transmitting tract, tissue sections were stained with a solution of 0.5% Alcian Blue and counterstained with 0.5% Neutral Red as previously described (Zúñiga-Mayo et al., 2012). A coverslip was placed over the sample and pictures were taken using a DM6000B microscope (Leica).

Pollen tube growth was analyzed using Aniline Blue staining as described by (Jiang et al., 2005). Both wildtype and mutant flowers were emasculated at the anthesis stage and 24 h later were hand-pollinated. The next day the gynoecia were collected and fixed in ethanol:acetic acid (3:1) for 2 h at room temperature. The fixed gynoecia were washed three times with distilled water and treated in softening solution of 8 M NaOH overnight. Then, the gynoecia were washed three times in distilled water and stained in aniline blue solution (0.1% aniline blue in 150 mM K_2_HPO_4_ buffer, pH 11) for 5 h in the dark. Gynoecia were observed and photographed with a DM6000B fluorescence microscope under UV light (Leica).

### Accession numbers

Sequence data from this article can be found in the European Nucleotide Archive (ENA) under the accession number XXXXX.

## Acknowledgments

We thank Joanna Serwatowska and Katarzyna Oktaba from CINVESTAV-IPN, Unidad Irapuato for technical advice on library preparations. V.L.G. and J.J.B.G. were supported by the Consejo Nacional de Humanidades, Ciencias y Tecnologías (CONAHCYT, Mexico) with a Ph.D. fellowship (487657 and 783880, respectively). The work in the S.d.F. laboratory was financed by the CONAHCYT grants CB-2012-177739, FC-2015-2/1061 and CB-2017-2018-A1-S-10126. S.d.F. is grateful for the participation in the European Union projects H2020-MSCA-RISE-2020 EVOfruland (101007738) and H2020-MSCA-RISE-2019 MAD (872417). N.J.P. thanks support of Faculty of Arts & Sciences, University of Toronto. M.R.P. thanks support of the ERASMUS program.

## Author contributions

Conceptualization, V.L.G., S.d.F.; Investigation, V.L.G. (LAM, library preparations, RNA-seq analysis, genetics, histology), J.J.B.G. (promoter analysis); Database: M.R.P., A.P., N.J.P.; Writing – Original draft, V.L.G.; Writing – Reviewing and editing, V.L.G., S.d.F.; Funding Acquisition, S.d.F. All authors have read and agreed to the published version of the manuscript.

## Declaration of interests

The authors declare no competing interests.

## Supplemental legends

**Supplemental Figure 1. Overview of the Laser Assisted Microdissection and RNA-Seq process.**

**Supplemental Figure 2. Principal component analysis (PCA).**

**Supplemental Figure 3. Circle plots with the unique transcription factors of the DEGs of each tissue.**

**Supplemental Figure 4. Functional analyses of candidate genes.**

**Supplemental Figure 5. The medial domain transcriptomic profile is shared with YFPP sample.**

**Supplemental Table S1. Tissues LAM, quality of RNA samples used to make the libraries and results of the alignment.**

**Supplemental Table S2. Gene expression without cut off value for each tissue.**

**Supplemental Table S3. Gene expression in CMM.**

**Supplemental Table S4. Gene expression in PC.**

**Supplemental Table S5. Gene expression in SEP.**

**Supplemental Table S6. Gene expression in VV.**

**Supplemental Table S7. Clustering of gene expression profiles.**

**Supplemental Table S8. Tissue-specific expressed genes in CMM, PC, SEP, CMM.**

**Supplemental Table S9. Tissue-specific transcription factors expressed in CMM, PC, SEP, VV.**

**Supplemental Table S10. Expression of tissue-specific genes.**

**Supplemental Table S11. Transcription factors expressed in CMM, PC, SEP, VV.**

**Supplemental Table S12. Expression profiles of the 86 genes (without the cut-off value 4 TPM) reported by Reyes-Olalde et al., 2017, in each tissue.**

**Supplemental Table S13. Transcription factors expressed in each tissue.**

**Supplemental Table S14. Dynamics of auxin pathway genes in each tissue.**

**Supplemental Table S15. Dynamics of cytokinin pathway genes in each tissue.**

**Supplemental Table S16. Cell cycle core gene expression in each tissue.**

**Supplemental Table S17. Differentially expressed genes (DEGs) of CMM.**

**Supplemental Table S18. Differentially expressed genes (DEGs) of PC.**

**Supplemental Table S19. Differentially expressed genes (DEGs) of SEP.**

**Supplemental Table S20. Differentially expressed genes (DEGs) of VV.**

**Supplemental Table S21. DEGs encoding unique transcription factors in the CMM, PC, SEP, VV.**

**Supplemental Table S22. Genes previously reported to be expressed and to participate in medial and lateral domain development.**

**Supplemental Table S23. Results of the GO analysis of enriched DEGs of CMM, PC, SEP, VV.**

**Supplemental Table S24. DEG selected for functional analysis.**

## Notes

### Competing Interest Statement

The authors have declared no competing interest.

## References

Alvarez, J., & Smyth, D. R. (1999). CRABS CLAW and SPATULA, two Arabidopsis genes that control carpel development in parallel with AGAMOUS. Development, 126(11), 2377–2386. https://doi.org/10.1242/dev.126.11.2377

Andersen, S. U., Buechel, S., Zhao, Z., Ljung, K., Novák, O., Busch, W., Schuster, C., & Lohmann, J. U. (2008). Requirement of B2-type cyclin-dependent kinases for meristem integrity in Arabidopsis thaliana. Plant Cell, 20(1). https://doi.org/10.1105/tpc.107.054676

Andrews, S. (2010). FastQC - A quality control tool for high throughput sequence data. http://www.bioinformatics.babraham.ac.uk/projects/fastqc/. Babraham Bioinformatics.

Bailey, T. L., & Grant, C. E. (2021). SEA: Simple Enrichment Analysis of motifs. BioRxiv.

Balanzá, V., Navarrete, M., Trigueros, M., & Ferrándiz, C. (2006). Patterning the female side of Arabidopsis: the importance of hormones. In Journal of Experimental Botany (Vol. 57, Issue 13). https://doi.org/10.1093/jxb/erl188

Belmonte, M. F., Kirkbride, R. C., Stone, S. L., Pelletier, J. M., Bui, A. Q., Yeung, E. C., Hashimoto, M., Fei, J., Harada, C. M., Munoz, M. D., Le, B. H., Drews, G. N., Brady, S. M., Goldberg, R. B., & Harada, J. J. (2013). Comprehensive developmental profiles of gene activity in regions and subregions of the Arabidopsis seed. Proceedings of the National Academy of Sciences of the United States of America, 110(5). https://doi.org/10.1073/pnas.1222061110

Bolger, A. M., Lohse, M., & Usadel, B. (2014). Trimmomatic: A flexible trimmer for Illumina sequence data. Bioinformatics, 30(15). https://doi.org/10.1093/bioinformatics/btu170

Bonnot, T., Gillard, M., & Nagel, D. (2019). A Simple Protocol for Informative Visualization of Enriched Gene Ontology Terms. BIO-PROTOCOL, 9(22). https://doi.org/10.21769/bioprotoc.3429

Boudolf, V., Rombauts, S., Naudts, M., Inzé, D., & De Veylder, L. (2001). Identification of novel cyclin-dependent kinases interacting with the CKS1 protein of Arabidopsis. Journal of Experimental Botany, 52(359). https://doi.org/10.1093/jxb/52.359.1381

Bowman, J. L., Baum, S. F., Eshed, Y., Putterill, J., & Alvarez, J. (1999). Molecular genetics of gynoecium development in Arabidopsis. In Current topics in developmental biology (Vol. 45). https://doi.org/10.1016/s0070-2153(08)60316-6

Bowman, J. L., & Smyth, D. R. (1999). CRABS CLAW, a gene that regulates carpel and nectary development in Arabidopsis, encodes a novel protein with zinc finger and helix-loop-helix domains. Development, 126(11). https://doi.org/10.1242/dev.126.11.2387

Bray, N. L., Pimentel, H., Melsted, P., & Pachter, L. (2016). Near-optimal probabilistic RNA-seq quantification. Nature Biotechnology, 34(5). https://doi.org/10.1038/nbt.3519

Carter, A. D., Bonyadi, R., & Gifford, M. L. (2013). The use of fluorescence-activated cell sorting in studying plant development and environmental responses. In International Journal of Developmental Biology (Vol. 57, Issues 6–8). https://doi.org/10.1387/ijdb.130195mg

Cervantes-Pérez, S. A., Thibivillliers, S., Tennant, S., & Libault, M. (2022). Review: Challenges and perspectives in applying single nuclei RNA-seq technology in plant biology. In Plant Science (Vol. 325). https://doi.org/10.1016/j.plantsci.2022.111486

Chávez Montes, R. A., Serwatowska, J., & De Folter, S. (2016). Laser-assisted microdissection to study global transcriptional changes during plant embryogenesis. In Somatic Embryogenesis: Fundamental Aspects and Applications. https://doi.org/10.1007/978-3-319-33705-0_27

Chiu, W. H., Chandler, J., Cnops, G., Van Lijsebettens, M., & Werr, W. (2007). Mutations in the TORNADO2 gene affect cellular decisions in the peripheral zone of the shoot apical meristem of Arabidopsis thaliana. Plant Molecular Biology, 63(6). https://doi.org/10.1007/s11103-006-9105-z

Crawford, B. C. W., Ditta, G., & Yanofsky, M. F. (2007). The NTT Gene Is Required for Transmitting-Tract Development in Carpels of Arabidopsis thaliana. Current Biology, 17(13). https://doi.org/10.1016/j.cub.2007.05.079

Crawford, B. C. W., & Yanofsky, M. F. (2011). Half filled promotes reproductive tract development and fertilization efficiency in Arabidopsis thaliana. Development, 138(14). https://doi.org/10.1242/dev.067793

Cubas, P., Lauter, N., Doebley, J., & Coen, E. (1999). The TCP domain: A motif found in proteins regulating plant growth and development. Plant Journal, 18(2). https://doi.org/10.1046/j.1365-313X.1999.00444.x

Day, R. C., Grossniklaus, U., & Macknight, R. C. (2005). Be more specific! Laser-assisted microdissection of plant cells. In Trends in Plant Science (Vol. 10, Issue 8). https://doi.org/10.1016/j.tplants.2005.06.006

Deal, R. B., & Henikoff, S. (2011). The INTACT method for cell typeg-specific gene expression and chromatin profiling in Arabidopsis thaliana. Nature Protocols, 6(1). https://doi.org/10.1038/nprot.2010.175

Deb, J., Bland, H. M., & Østergaard, L. (2018). Developmental cartography: coordination via hormonal and genetic interactions during gynoecium formation. In Current Opinion in Plant Biology (Vol. 41). https://doi.org/10.1016/j.pbi.2017.09.004

Denyer, T., & Timmermans, M. C. P. (2022). Crafting a blueprint for single-cell RNA sequencing. In Trends in Plant Science (Vol. 27, Issue 1). https://doi.org/10.1016/j.tplants.2021.08.016

Di Marzo, M., Roig-Villanova, I., Zanchetti, E., Caselli, F., Gregis, V., Bardetti, P., Chiara, M., Guazzotti, A., Caporali, E., Mendes, M. A., Colombo, L., & Kater, M. M. (2020). MADS-box and bHLH transcription factors coordinate transmitting tract development in arabidopsis thaliana. Frontiers in Plant Science, 11. https://doi.org/10.3389/fpls.2020.00526

Dinneny, J. R., Weigel, D., & Yanofsky, M. F. (2005). A genetic framework for fruit patterning in Arabidopsis thaliana. Development, 132(21). https://doi.org/10.1242/dev.02062

Ferrándiz, C., Fourquin, C., Prunet, N., Scutt, C. P., Sundberg, E., Trehin, C., & Vialette-Guiraud, A. C. M. (2010). Chapter 1 – Carpel Development. In Advances in Botanical Research (Vol. 55).

Florez Rueda, A. M., Grossniklaus, U., & Schmidt, A. (2016). Laser-assisted microdissection (LAM) as a tool for transcriptional profiling of individual cell types. Journal of Visualized Experiments, 2016(111). https://doi.org/10.3791/53916

Foreman, J., White, J. N., Graham, I. A., Halliday, K. J., & Josse, E. marie. (2011). Shedding light on flower development: Phytochrome B regulates gynoecium formation in association with the transcription factor SPATULA. Plant Signaling and Behavior, 6(4). https://doi.org/10.4161/psb.6.4.14496

Franco-Zorrilla, J. M., Cubas, P., Jarillo, J. A., Fernández-Calvín, B., Salinas, J., & Martínez-Zapater, J. M. (2002). AtREM1, a member of a new family of B3 domain-containing genes, is preferentially expressed in reproductive meristems. Plant Physiology, 128(2). https://doi.org/10.1104/pp.010323

Franco-Zorrilla, J. M., López-Vidriero, I., Carrasco, J. L., Godoy, M., Vera, P., & Solano, R. (2014). DNA-binding specificities of plant transcription factors and their potential to define target genes. Proceedings of the National Academy of Sciences of the United States of America, 111(6). https://doi.org/10.1073/pnas.1316278111

Girin, T., Paicu, T., Stephenson, P., Fuentes, S., Körner, E., O’Brien, M., Sorefan, K., Wood, T. A., Balanzá, V., Ferrándiz, C., Smyth, D. R., & Østergaard, L. (2011). INDEHISCENT and SPATULA interact to specify carpel and valve margin tissue and thus promote seed dispersal in Arabidopsis. Plant Cell, 23(10). https://doi.org/10.1105/tpc.111.090944

Gremski, K., Ditta, G., & Yanofsky, M. F. (2007). The HECATE genes regulate female reproductive tract development in Arabidopsis thaliana. Development, 134(20). https://doi.org/10.1242/dev.011510

Groszmann, M., Paicu, T., & Smyth, D. R. (2008). Functional domains of SPATULA, a bHLH transcription factor involved in carpel and fruit development in Arabidopsis. Plant Journal, 55(1). https://doi.org/10.1111/j.1365-313X.2008.03469.x

Gu, Q., Ferrándiz, C., Yanofsky, M. F., & Martienssen, R. (1998). The FRUITFULL MADS-box gene mediates cell differentiation during Arabidopsis fruit development. Development, 125(8). https://doi.org/10.1242/dev.125.8.1509

Gutierrez, C. (2009). The Arabidopsis Cell Division Cycle. The Arabidopsis Book, 7. https://doi.org/10.1199/tab.0120

Heisler, M. G. B., Atkinson, A., Bylstra, Y. H., Walsh, R., & Smyth, D. R. (2001). SPATULA, a gene that controls development of carpel margin tissues in Arabidopsis, encodes a bHLH protein. Development, 128(7). https://doi.org/10.1242/dev.128.7.1089

Herrera-Ubaldo, H., Campos, S. E., López-Gómez, P., Luna-García, V., Zúñiga-Mayo, V. M., Armas-Caballero, G. E., González-Aguilera, K. L., DeLuna, A., Marsch-Martínez, N., Espinosa-Soto, C., & de Folter, S. (2023). The protein–protein interaction landscape of transcription factors during gynoecium development in Arabidopsis. Molecular Plant, 16(1). https://doi.org/10.1016/j.molp.2022.09.004

Herrera-Ubaldo, H., & de Folter, S. (2022). Gynoecium and fruit development in Arabidopsis. In Development (Cambridge) (Vol. 149, Issue 5). https://doi.org/10.1242/dev.200120

Herrera-Ubaldo, H., Lozano-Sotomayor, P., Ezquer, I., Di Marzo, M., Montes, R. A. C., Goḿez-Felipe, A., Pablo-Villa, J., Diaz-Ramirez, D., Ballester, P., Ferrańdiz, C., Sagasser, M., Colombo, L., Marsch-Martıńez, N., & de Folter, S. (2019). New roles of NO TRANSMITTING TRACT and SEEDSTICK during medial domain development in arabidopsis fruits. Development (Cambridge), 146(1). https://doi.org/10.1242/dev.172395

Inzé, D., & De Veylder, L. (2006). Cell cycle regulation in plant development. In Annual Review of Genetics (Vol. 40). https://doi.org/10.1146/annurev.genet.40.110405.090431

Jiang, L., Yang, S. L., Xie, L. F., Puah, C. S., Zhang, X. Q., Yang, W. C., Sundaresan, V., & Ye, D. (2005). VANGUARD1 encodes a pectin methylesterase that enhances pollen tube growth in the Arabidopsis style and transmitting tract. Plant Cell, 17(2). https://doi.org/10.1105/tpc.104.027631

Kamiuchi, Y., Yamamoto, K., Furutani, M., Tasaka, M., & Aida, M. (2014). The CUC1 and CUC2 genes promote carpel margin meristem formation during Arabidopsis gynoecium development. Frontiers in Plant Science, 5(APR). https://doi.org/10.3389/fpls.2014.00165

Kerk, N. M., Ceserani, T., Lorraine Tausta, S., Sussex, I. M., & Nelson, T. M. (2003). Laser capture microdissection of cells from plant tissues. Plant Physiology, 132(1). https://doi.org/10.1104/pp.102.018127

Larsson, E., Franks, R. G., & Sundberg, E. (2013). Auxin and the Arabidopsis thaliana gynoecium. In Journal of Experimental Botany (Vol. 64, Issue 9). https://doi.org/10.1093/jxb/ert099

Larsson, E., Roberts, C. J., Claes, A. R., Franks, R. G., & Sundberg, E. (2014). Polar auxin transport is essential for medial versus lateral tissue specification and vascular-mediated valve outgrowth in Arabidopsis gynoecia. Plant Physiology, 166(4). https://doi.org/10.1104/pp.114.245951

Liljegren, S. J., Roeder, A. H. K., Kempin, S. A., Gremski, K., Østergaard, L., Guimil, S., Reyes, D. K., & Yanofsky, M. F. (2004). Control of fruit patterning in Arabidopsis by INDEHISCENT. Cell, 116(6). https://doi.org/10.1016/S0092-8674(04)00217-X

Liu, J., Sheng, L., Xu, Y., Li, J., Yang, Z., Huang, H., & Xu, L. (2014). WOX11 and 12 are involved in the first-step cell fate transition during de novo root organogenesis in Arabidopsis. Plant Cell, 26(3). https://doi.org/10.1105/tpc.114.122887

Lucero, L. E., Uberti-Manassero, N. G., Arce, A. L., Colombatti, F., Alemano, S. G., & Gonzalez, D. H. (2015). TCP15 modulates cytokinin and auxin responses during gynoecium development in Arabidopsis. Plant Journal, 84(2). https://doi.org/10.1111/tpj.12992

Mantegazza, O., Gregis, V., Mendes, M. A., Morandini, P., Alves-Ferreira, M., Patreze, C. M., Nardeli, S. M., Kater, M. M., & Colombo, L. (2014). Analysis of the Arabidopsis REM gene family predicts functions during flower development. In Annals of Botany (Vol. 114, Issue 7). https://doi.org/10.1093/aob/mcu124

Marsch-Martínez, N., & de Folter, S. (2016). Hormonal control of the development of the gynoecium. In Current Opinion in Plant Biology (Vol. 29). https://doi.org/10.1016/j.pbi.2015.12.006

Marsch-Martinez, N., Reyes-Olalde, J. I., Ramos-Cruz, D., Lozano-Sotomayor, P., Zúñiga-Mayo, V. M., & de Folter, S. (2012a). Hormones talking: does hormonal cross-talk shape the Arabidopsis gynoecium? Plant Signaling and Behavior, 7(12). https://doi.org/10.4161/psb.22422

Marsch-Martínez, N., Ramos-Cruz, D., Irepan Reyes-Olalde, J., Lozano-Sotomayor, P., Zúñiga-Mayo, V. M., & de Folter, S. (2012b). The role of cytokinin during Arabidopsis gynoecia and fruit morphogenesis and patterning. Plant Journal, 72(2). https://doi.org/10.1111/j.1365-313X.2012.05062.x

Martin, M. (2011). Cutadapt removes adapter sequences from high-throughput sequencing reads. EMBnet.Journal, 17(1). https://doi.org/10.14806/ej.17.1.200

Matsumoto, N., & Okada, K. (2001). A homeobox gene, PRESSED FLOWER, regulates lateral axis-dependent development of Arabidopsis flowers. Genes and Development, 15(24). https://doi.org/10.1101/gad.931001

Menges, M., De Jager, S. M., Gruissem, W., & Murray, J. A. H. (2005). Global analysis of the core cell cycle regulators of Arabidopsis identifies novel genes, reveals multiple and highly specific profiles of expression and provides a coherent model for plant cell cycle control. Plant Journal, 41(4). https://doi.org/10.1111/j.1365-313X.2004.02319.x

Moubayidin, L., & Østergaard, L. (2014). Dynamic Control of Auxin Distribution Imposes a Bilateral-to-Radial Symmetry Switch during Gynoecium Development. Current Biology, 24(22). https://doi.org/10.1016/j.cub.2014.09.080

Moubayidin, L., & Østergaard, L. (2017). Gynoecium formation: an intimate and complicated relationship. In Current Opinion in Genetics and Development (Vol. 45). https://doi.org/10.1016/j.gde.2017.02.005

Müller, C. J., Larsson, E., Spíchal, L., & Sundberg, E. (2017). Cytokinin-auxin crosstalk in the gynoecial primordium ensures correct domain patterning. Plant Physiology, 175(3). https://doi.org/10.1104/pp.17.00805

Nahar, M. A. U., Ishida, T., Smyth, D. R., Tasaka, M., & Aida, M. (2012). Interactions of CUP-SHAPED COTYLEDON and SPATULA genes control carpel margin development in arabidopsis thaliana. Plant and Cell Physiology, 53(6). https://doi.org/10.1093/pcp/pcs057

Nemhauser, J. L., Feldman, L. J., & Zambryski, P. C. (2000). Auxin and ETTIN in Arabidopsis gynoecium morphogenesis. Development, 127(18). https://doi.org/10.1242/dev.127.18.3877

Panikashvili, D., Shi, J. X., Schreiber, L., & Aharoni, A. (2011). The Arabidopsis ABCG13 transporter is required for flower cuticle secretion and patterning of the petal epidermis. New Phytologist, 190(1). https://doi.org/10.1111/j.1469-8137.2010.03608.x

Pekker, I., Alvarez, J. P., & Eshed, Y. (2005). Auxin response factors mediate Arabidopsis organ asymmetry via modulation of KANADI activity. Plant Cell, 17(11). https://doi.org/10.1105/tpc.105.034876

Peterson, R., Slovin, J. P., & Chen, C. (2010). A simplified method for differential staining of aborted and non-aborted pollen grains. International Journal of Plant Biology, 1(2). https://doi.org/10.4081/pb.2010.e13

Rajani, S., & Sundaresan, V. (2001). The Arabidopsis myc/bHLH gene alcatraz enables cell separation in fruit dehiscence. Current Biology, 11(24). https://doi.org/10.1016/S0960-9822(01)00593-0

Ramos-Pulido, J., & de Folter, S. (2023). Organogenic events during gynoecium and fruit development in Arabidopsis. In Current Opinion in Plant Biology, in press.

Reyes-Olalde, J. I., & de Folter, S. (2019). Control of stem cell activity in the carpel margin meristem (CMM) in Arabidopsis. In Plant Reproduction (Vol. 32, Issue 2). https://doi.org/10.1007/s00497-018-00359-0

Reyes-Olalde, J. I., Zuñiga-Mayo, V. M., Chávez Montes, R. A., Marsch-Martínez, N., & de Folter, S. (2013). Inside the gynoecium: At the carpel margin. In Trends in Plant Science (Vol. 18, Issue 11). https://doi.org/10.1016/j.tplants.2013.08.002

Reyes-Olalde, J. I., Zúñiga-Mayo, V. M., Serwatowska, J., Chavez Montes, R. A., Lozano-Sotomayor, P., Herrera-Ubaldo, H., Gonzalez-Aguilera, K. L., Ballester, P., Ripoll, J. J., Ezquer, I., Paolo, D., Heyl, A., Colombo, L., Yanofsky, M. F., Ferrandiz, C., Marsch-Martínez, N., & de Folter, S. (2017). The bHLH transcription factor SPATULA enables cytokinin signaling, and both activate auxin biosynthesis and transport genes at the medial domain of the gynoecium. PLoS Genetics, 13(4). https://doi.org/10.1371/journal.pgen.1006726

Reymond, M. C., Brunoud, G., Chauvet, A., Martínez-Garcia, J. F., Martin-Magniette, M. L., Monéger, F., & Scutt, C. P. (2012). A Light-Regulated Genetic Module Was Recruited to Carpel Development in Arabidopsis following a Structural Change to SPATULA. Plant Cell, 24(7). https://doi.org/10.1105/tpc.112.097915

Rich-Griffin, C., Stechemesser, A., Finch, J., Lucas, E., Ott, S., & Schäfer, P. (2020). Single-Cell Transcriptomics: A High-Resolution Avenue for Plant Functional Genomics. In Trends in Plant Science (Vol. 25, Issue 2). https://doi.org/10.1016/j.tplants.2019.10.008

Robert, H. S., Crhak Khaitova, L., Mroue, S., & Benková, E. (2015). The importance of localized auxin production for morphogenesis of reproductive organs and embryos in Arabidopsis. Journal of Experimental Botany, 66(16). https://doi.org/10.1093/jxb/erv256

Robinson, M. D., McCarthy, D. J., & Smyth, G. K. (2009). edgeR: A Bioconductor package for differential expression analysis of digital gene expression data. Bioinformatics, 26(1). https://doi.org/10.1093/bioinformatics/btp616

Roeder, A. H. K., & Yanofsky, M. F. (2006). Fruit Development in Arabidopsis. The Arabidopsis Book, 4, e0075. https://doi.org/10.1199/tab.0075

Schaller, G. E., Bishopp, A., & Kieber, J. J. (2015). The yin-yang of hormones: Cytokinin and auxin interactions in plant development. Plant Cell, 27(1), 44–63. https://doi.org/10.1105/tpc.114.133595

Sehra, B., & Franks, R. G. (2015). Auxin and cytokinin act during gynoecial patterning and the development of ovules from the meristematic medial domain. Wiley Interdiscip Rev Dev Biol, (Vol. 4). http://dx.doi.org/10.1002/wdev.1193.

Shimotohno, A., Aki, S. S., Takahashi, N., & Umeda, M. (2021). Regulation of the Plant Cell Cycle in Response to Hormones and the Environment. In Annual Review of Plant Biology (Vol. 72). https://doi.org/10.1146/annurev-arplant-080720-103739

Simonini, S., & Østergaard, L. (2019). Female reproductive organ formation: A multitasking endeavor. In Current Topics in Developmental Biology (Vol. 131). https://doi.org/10.1016/bs.ctdb.2018.10.004

Smyth, D. R., Bowman, J. L., & Meyerowitz, E. M. (1990). Early flower development in Arabidopsis. Plant Cell, 2(8). https://doi.org/10.1105/tpc.2.8.755

Soneson, C., Love, M. I., & Robinson, M. D. (2015). Differential analyses for RNA-seq: transcript-level estimates improve gene-level inferences. F1000Research, 4. https://doi.org/10.12688/f1000research.7563.1

Supek, F., Bošnjak, M., Škunca, N., & Šmuc, T. (2011). Revigo summarizes and visualizes long lists of gene ontology terms. PLoS ONE, 6(7). https://doi.org/10.1371/journal.pone.0021800

Takeda, S., Iwasaki, A., Tatematsu, K., & Okada, K. (2014). The half-size ABC transporter FOLDED PETALS 2/ABCG13 is involved in petal elongation through narrow spaces in Arabidopsis thaliana floral buds. Plants, 3(3). https://doi.org/10.3390/plants3030348

Taylor-Teeples, M., Ron, M., & Brady, S. M. (2011). Novel biological insights revealed from cell type-specific expression profiling. In Current Opinion in Plant Biology (Vol. 14, Issue 5). https://doi.org/10.1016/j.pbi.2011.05.007

Vandepoele, K., Raes, J., De Veylder, L., Rouzé, P., Rombauts, S., & Inzé, D. (2002). Genome-wide analysis of core cell cycle genes in Arabidopsis. Plant Cell, 14(4). https://doi.org/10.1105/tpc.010445

Villarino, G. H., Hu, Q., Manrique, S., Flores-Vergara, M., Sehra, B., Robles, L., Brumos, J., Stepanova, A. N., Colombo, L., Sundberg, E., Heber, S., & Franks, R. G. (2016). Transcriptomic signature of the SHATTERPROOF2 expression domain reveals the meristematic nature of Arabidopsis gynoecial medial domain. Plant Physiology, 171(1). https://doi.org/10.1104/pp.15.01845

Winter, D., Vinegar, B., Nahal, H., Ammar, R., Wilson, G. V., & Provart, N. J. (2007). An “electronic fluorescent pictograph” Browser for exploring and analyzing large-scale biological data sets. PLoS ONE, 2(8). https://doi.org/10.1371/journal.pone.0000718

Wuest, S. E., & Grossniklaus, U. (2014). Laser-assisted microdissection applied to floral tissues. Methods in Molecular Biology, 1110. https://doi.org/10.1007/978-1-4614-9408-9_19

Wynn, A. N., Rueschhoff, E. E., & Franks, R. G. (2011). Transcriptomic characterization of a synergistic genetic interaction during carpel margin meristem development in arabidopsis thaliana. PLoS ONE, 6(10). https://doi.org/10.1371/journal.pone.0026231

Zanetti, M. E., Chang, I. F., Gong, F., Galbraith, D. W., & Bailey-Serres, J. (2005). Immunopurification of polyribosomal complexes of Arabidopsis for global analysis of gene expression. In Plant Physiology (Vol. 138, Issue 2). https://doi.org/10.1104/pp.105.059477

Zheng, Y., Jiao, C., Sun, H., Rosli, H. G., Pombo, M. A., Zhang, P., Banf, M., Dai, X., Martin, G. B., Giovannoni, J. J., Zhao, P. X., Rhee, S. Y., & Fei, Z. (2016). iTAK: A Program for Genome-wide Prediction and Classification of Plant Transcription Factors, Transcriptional Regulators, and Protein Kinases. In Molecular Plant (Vol. 9, Issue 12). https://doi.org/10.1016/j.molp.2016.09.014

Zúñiga-Mayo, V. M., Gómez-Felipe, A., Herrera-Ubaldo, H., & de Folter, S. (2019). Gynoecium development: Networks in Arabidopsis and beyond. In Journal of Experimental Botany (Vol. 70, Issue 5). https://doi.org/10.1093/jxb/erz026

Zúñiga-Mayo, V. M., Marsch-Martínez, N., & de Folter, S. (2012). JAIBA, a class-II HD-ZIP transcription factor involved in the regulation of meristematic activity, and important for correct gynoecium and fruit development in Arabidopsis. Plant Journal, 71(2). https://doi.org/10.1111/j.1365-313X.2012.04990.x

